# Interaction of the *Xanthomonas* effectors XopQ and XopX results in induction of rice immune responses

**DOI:** 10.1101/2020.02.07.938365

**Authors:** Sohini Deb, Palash Ghosh, Hitendra K. Patel, Ramesh V. Sonti

## Abstract

*Xanthomonas oryzae* pv. *oryzae* uses several type III secretion system (T3SS) effectors, namely XopN, XopQ, XopX, and XopZ, to suppress rice immune responses that are induced following treatment with cell wall degrading enzymes. Here we show that the T3SS secreted effector XopX interacts with two of the eight rice 14-3-3 proteins. Mutants of XopX that are defective in 14-3-3 binding are also defective in suppression of immune responses, suggesting that interaction with 14-3-3 proteins is required for suppression of host innate immunity. However, *Agrobacterium* mediated delivery of both XopX and XopQ into rice cells results in induction of rice immune responses. These immune responses are not observed when either protein is individually delivered into rice cells. XopQ-XopX induced rice immune responses are not observed in a XopX mutant that is defective in 14-3-3 binding. Yeast two-hybrid and BiFC assays indicate that XopQ and XopX interact with each other. In a screen for *Xanthomonas* effectors which can suppress XopQ-XopX induced rice immune responses, five effectors were identified, namely XopU, XopV, XopP, XopG and AvrBs2, which were able to do so. These results suggest a complex interplay of *Xanthomonas* T3SS effectors in suppression of pathogen triggered immunity and effector triggered immunity to promote virulence on rice.

**Significance statement:** This work studies the role of the type III effector XopX in the suppression and induction of rice immune responses, by differential interaction with the 14-3-3 proteins, or with the type III effector XopQ respectively. We have also identified a subset of type III effectors which can suppress this form of immune responses.

## Introduction

Plants are subject to attack by various pathogenic organisms. To combat these pathogens, plants have evolved an immune system that is inducible and multi-layered. The first line of defence arises from the recognition of signature patterns on the pathogen, known as pattern - associated molecular patterns (PAMPs) through specific pattern recognition receptors (PRRs) at the plant cell surface (Saijo et al., 2018). This leads to activation of PAMP-triggered immunity (PTI), which is characterised by production of reactive oxygen species (ROS), callose deposition and expression of defence-related genes (Jones and Dangl, 2006). Successful pathogens suppress this immune response with the help of effectors, which in Gram-negative bacteria are primarily secreted through their type III secretion system. For this, they target specific plant proteins involved in the immune response cascade (White and Yang, 2009). For example, the *Xanthomonas oryzae* pv. *oryzae* effector XopQ interacts with the rice 14-3-3 proteins to suppress immune responses (Deb et al., 2019). The AvrPto effector has also been shown to interact with the plant receptor FLS2, which recognizes the bacterial flagellin, further suppressing host PTI (Xiang et al., 2008). The XopJ effector of *Xanthomonas campestris* mediates inhibition of the proteasome to interfere with SA-dependent defence response (Üstün et al., 2013). The second layer of plant defence involves recognition of these effectors by the plant, via specific disease resistance genes (R genes), which are usually nucleotide-binding leucine-rich repeat (NB-LRR) proteins, leading to effector-triggered immunity (ETI) (Chisholm et al., 2006). ETI generally leads to a strong defence reaction called the hypersensitive response (HR), characterized by rapid cell death and local necrosis, which prevents further spread of pathogen (Oh and Martin, 2011).

The type III effectors Xanthomonas outer protein Q (XopQ) and XopX of *X. oryzae* pv. *oryzae* had been identified as two out of four type III effectors which could suppress the innate immune responses induced by treatment of a cell-wall degrading enzyme in rice (Sinha et al., 2013). The XopQ protein is highly conserved in Xanthomonads (Hajri et al., 2009, Moreira et al., 2010, Jalan et al., 2013, Potnis et al., 2011), and has been shown to require its biochemical activity for complete virulence in rice (Gupta et al., 2015). XopQ also has motifs for binding to 14-3-3 proteins (Dubrow et al., 2018, Deb et al., 2019). The 14-3-3 proteins are eukaryotic adapter proteins which play key roles in multiple cellular events in plants (Cotelle and Leonhardt, 2015, Cotelle et al., 2000). Rice has genes encoding for eight 14-3-3 isoforms, named as *gf14a, gf14b, gf14c, gf14d, gf14e, gf14f, gf14g* and *gf14h* (Chen et al., 2006).

A number of studies have shown the role of the 14-3-3 proteins in modulation of PTI and ETI (Lozano-Duran and Robatzek, 2015), by the binding of host 14-3-3 proteins with the type III effectors of the pathogen (Dubrow et al., 2018). The ability of *X. oryzae pv. oryzae* XopQ protein to interact with 14-3-3 proteins has been shown to be important for its ability to suppress rice immune responses (Deb et al., 2019). The *X. campestris* pv. *vesicatoria* XopQ has been shown to suppress effector triggered immunity (ETI)-associated cell death in pepper by interacting with the 14-3-3 protein TFT4 (Teper et al., 2014), and also suppresses cell death triggered by MAPKKKα (Teper et al., 2015). In two different studies, phosphorylation of the *Pseudomonas* ortholog, HopQ1, has been shown to enable binding of host 14-3-3 proteins, and has been shown to interact with the tomato 14-3-3 proteins TFT1 and TFT5 in a phosphorylation-dependent manner (Li et al., 2013, Giska et al., 2013). On the other hand, the *X. campestris* pv. *vesicatoria* XopX protein has been shown to contribute to virulence in pepper and tomato, and modulates the plant immune response, resulting in enhanced susceptibility of the plant (Metz et al., 2005). However, XopX also seems to be an inducer of rice immune responses wherein it was shown to weakly upregulate PTI marker genes (Stork et al., 2015). The *X. oryzae* pv. *oryzae* XopX protein has five putative 14-3-3 protein binding motifs.

In this study, we show that the *X. oryzae* pv. *oryzae* XopX protein interacts with rice 14-3-3 proteins and that this interaction is necessary for its ability to suppress rice immune responses. We also demonstrate that XopX interacts with XopQ. This interaction results in induction of rice immune responses in a 14-3-3 dependent manner. We show that several other *X. oryzae* pv. *oryzae* type III effectors such as XopU, XopV, XopP, XopG and AvrBs2 are able to suppress rice immune responses that are induced by XopQ-XopX. Overall, the results suggest that XopX can interact with a subset of rice 14-3-3 proteins as well as with the XopQ effector, and that this differential interaction leads to suppression or induction of immune responses respectively. Also, additional type III effectors of *X. oryzae* pv. *oryzae* may be involved in suppression of XopQ-XopX induced immune responses in rice.

## Results

### XopX interacts with two of the eight rice 14-3-3 proteins

Bioinformatic analysis revealed that *X. oryzae* pv. *oryzae* XopX has five putative 14-3-3 protein binding motifs encompassing the conserved residues serine-84 (mode-II motif ‘RASTSAP’; amino acid 80-86), serine-193 (mode-II motif ‘RGAISNP’; amino acid 189-195), threonine-430 (mode-II motif ‘RRDFTGP’; amino acid 426-432), serine-477 (mode-1 motif ‘RSESIP’; amino acid 474-479) and threonine-621 (mode-1 motif ‘RLFTGP’; amino acid 618-623). In order to check if XopX would interact with any of the rice 14-3-3 proteins, we cloned the wild-type *xopX* gene in the pDEST32 vector yielding the *BD∷xopX* (DNA-Binding Domain) clone (as listed in Supplementary table S2). We screened *xopX* against the eight rice 14-3-3 proteins cloned in the pDEST22 vector yielding the *AD∷gf14a-h* (Activation Domain) clones used in an earlier study (Deb et al., 2019) (listed in Supplementary table S2), using the yeast two-hybrid system and the yeast strain pJ694a (James et al., 1996). The one-to-one yeast two-hybrid screen indicated that the XopX protein showed physical interaction with two of the eight rice 14-3-3 proteins, Gf14d and Gf14e (Fig 1A). Growth of yeast on plates lacking leucine and tryptophan confirmed the presence of pDEST32*∷xopX* as well as pDEST22*∷gf14a-h* in the yeast strain. However, growth on plates lacking adenine, histidine, leucine and tryptophan, and supplemented with 3-amino-1,2,4-triazole (3-AT), indicated a positive interaction of XopX-Gf14d and XopX-Gf14e. We further tested this in a bimolecular fluorescence complementation (BiFC) assay, wherein *xopX* was cloned with the C-terminal of Venus fluorescent protein (VFP) yielding *cVFP∷xopX* and *gf14d* and *gf14e* were cloned with the N-terminal of VFP yielding *nVFP∷gf14d* and *nVFP∷gf14e* (as listed in Supplementary table S2). Infiltration of *Agrobacterium tumefaciens* AGL1 harbouring *cVFP∷xopX* and either *nVFP∷gf14d* or *nVFP∷gf14e* (as listed in Supplementary table S3) in *Nicotiana benthamiana* leaf epidermis revealed strong complementation of fluorescence for both Gf14d and Gf14e with XopX (Fig 1B).

**Fig 1.**
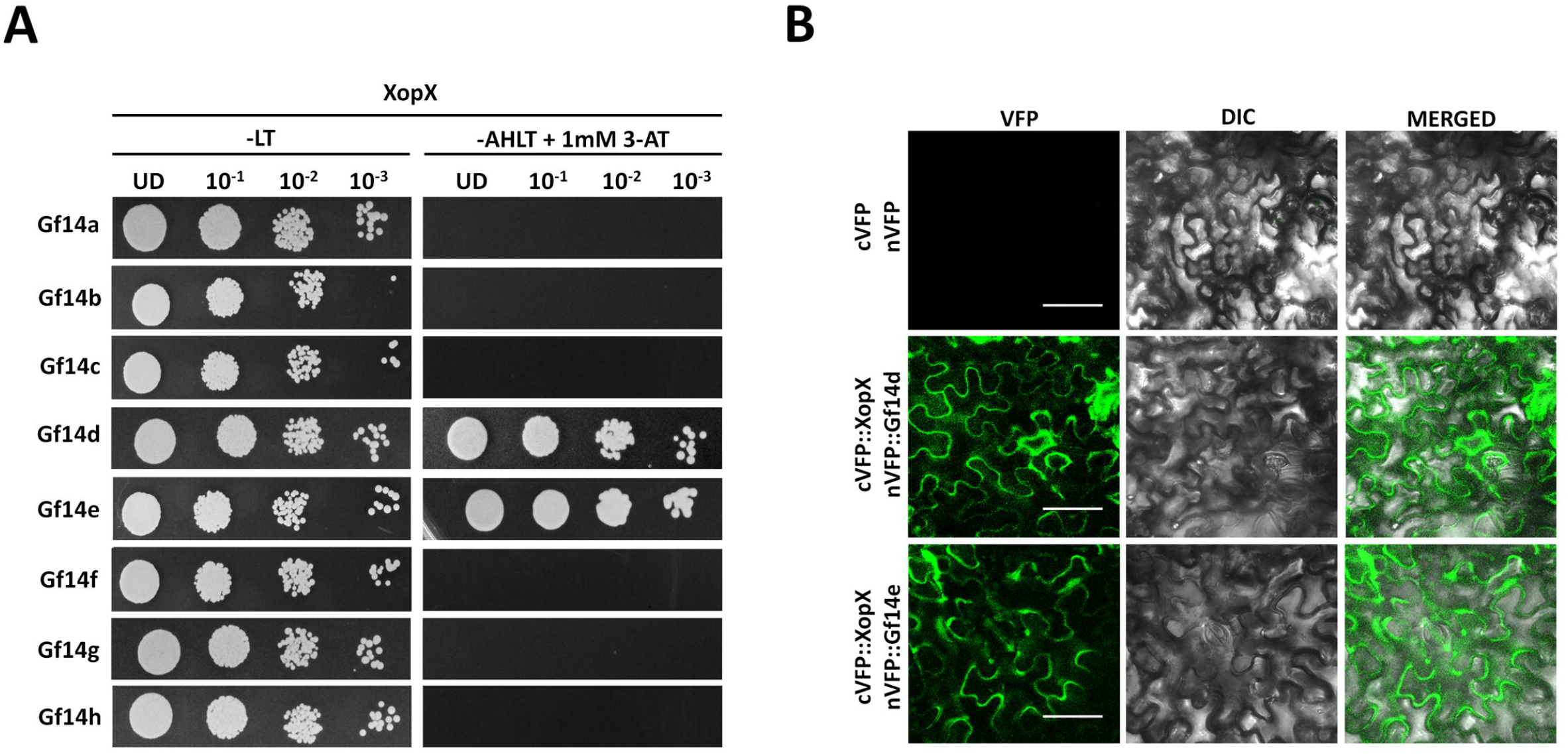
The *X. oryzae* pv. *oryzae* XopX protein interacts with two rice 14-3-3 proteins, Gf14d and Gf14e. **(A)** Yeast strain pJ694a containing pDEST32 vector expressing binding domain (BD) fusion with XopX was independently transformed with pDEST22 vector expressing activation domain (AD) fusion with Gf14a-h. Transformed colonies were serially diluted and spotted on the nonselective −LT (−Leu −Trp) double dropout (DDO) media and selective −AHLT (−Ade −Leu −Trp −His) quadruple dropout (QDO) media with 1mM 3-amino-1,2,4-triazole (3-AT). Observations were noted after 3 days of incubation at 30°C. Similar results were obtained in three independent experiments. **(B)** For BiFC analysis of XopX-14-3-3 interactions, leaves of *N. benthamiana* were syringe-infiltrated with a suspension of two *A. tumefaciens* AGL1 strains containing empty vectors alone or BiFC vectors expressing cVFP∷XopX and nVFP∷Gf14d or nVFP∷Gf14e. Fluorescence was visualised in a confocal microscope at 20x magnification and excitation wavelength (488nm) 48h after infiltration. Bar, 50μm. Similar results were obtained in three independent experiments.

### Mutation in the serine-193 and serine-477 containing 14-3-3 protein binding motifs of XopX abolishes its ability to interact with the rice 14-3-3 proteins

Since XopX showed interaction with rice 14-3-3 proteins, we further asked which of its 14-3-3 protein binding motifs was responsible for this interaction. In order to study this aspect, since 14-3-3 proteins are known to bind to client proteins in a phosphorylation-dependent manner, we individually mutated the conserved serine/ threonine residues in the five 14-3-3 protein binding motifs of XopX to alanine by site-directed mutagenesis. This yielded the phospho-null pENTR/D-TOPO constructs containing *xopX S84A, xopX S193A, xopX T430A, xopX S477A* and *xopX T621A*. These were then cloned into the yeast two-hybrid vector pDEST32 (Invitrogen) by Gateway® cloning (Invitrogen), yielding the clones *BD∷xopX S84A, BD∷xopX S193A, BD∷xopX T430A, BD∷xopX S477A* and *BD∷xopX T621A* (as listed in Supplementary table S2). These clones were further screened for interaction with pDEST22*∷gf14d* and pDEST22*∷gf14e* using the yeast two-hybrid system. It was observed that mutation of serine to alanine, individually, in motif-2 (*xopX S193A*) and motif-4 (*xopX S477A*), affected the ability of the XopX protein to interact with Gf14d as well as with Gf14e. When grown on selection plates for interaction, motif-2 XopX S193A was seen to lose interaction with both Gf14d and Gf14e (Fig 2A & B). However, motif-4 XopX S477A loses interaction with Gf14e but retains its ability to interact with Gf14d weakly (Fig 2A & B). However, *in-planta*, both the mutants XopX S193A and XopX S477A seem to lose interaction with both Gf14d (Fig 2C) and Gf14e (Fig 2D). The mutants in the other three 14-3-3 protein binding motifs, XopX S84A, XopX T430A and XopX T621A, seem to interact with both Gf14d and Gf14e as efficiently as wild-type XopX (Fig 2A & B). To further confirm this, phosphomimic mutants were made of the two motifs containing serine-193 and serine-477, yielding the XopX S193D and XopX S477D mutant proteins. XopX S193D as well as XopX S477D were seen to interact with both Gf14d and Gf14e, in yeast (Fig 2A & B) and *in-planta* (Fig 2C & D). Hence it appears that XopX requires phosphorylation of both motif-2 containing serine-193 and motif-4 containing serine-477 for interaction with Gf14d and Gf14e.

**Fig 2.**
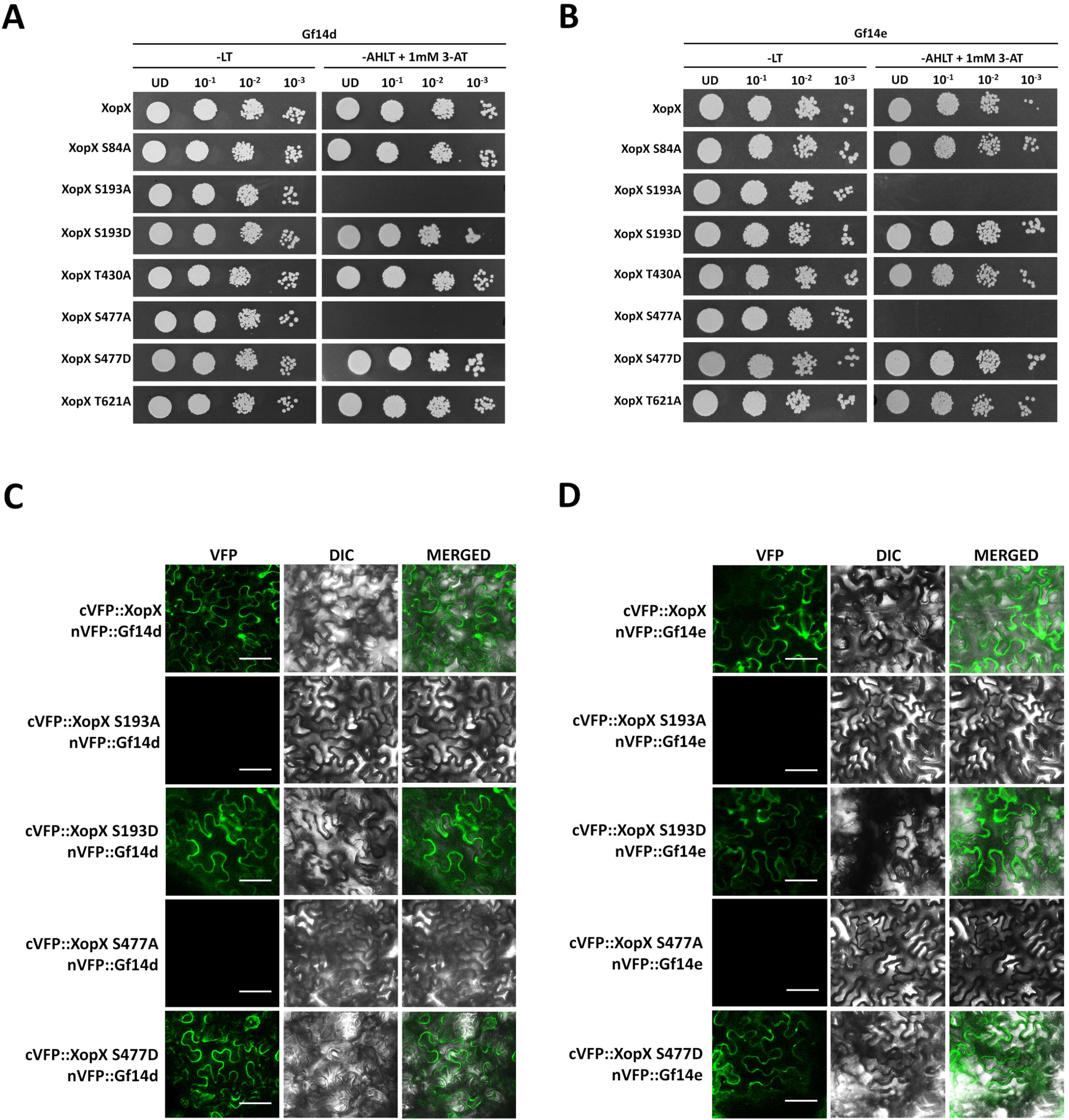
Mutation in the serine-193 and serine-477 containing 14-3-3 protein binding motifs of XopX abolishes its ability to interact with the 14-3-3 proteins Gf14d and Gf14e. **(A & B)** Yeast two-hybrid reporter strain pJ694a was transformed with the pDEST32 vector expressing binding domain (BD) fusion with XopX, XopX S84A, XopX S193A, XopX S193D, XopX T430A, XopX S477A, XopX S477D or XopX T621A and the prey vector pDEST22 vector expressing activation domain (AD) fusion with Gf14d **(A)** or Gf14e **(B)**. Transformed colonies were spotted on nonselective −LT (−Leucine −Tryptophan) double dropout (DDO) media and selective −AHLT (−Adenine −Histidine −Leucine −Tryptophan) quadruple dropout (QDO) media with 1mM 3-AT. Observations were noted after incubation at 30°C for 3 days. Similar results were obtained in three independent experiments. **(C & D)** BiFC analysis of XopX-14-3-3 interactions in *N. benthamiana*. Leaves were syringe-infiltrated with a suspension of two *A. tumefaciens* AGL1 strains expressing cVFP∷XopX, cVFP∷XopX S193A, cVFP∷XopX S193D, cVFP∷XopX S477A or cVFP∷XopX S477D and nVFP∷Gf14d or nVFP∷Gf14e. Fluorescence was visualised in a confocal microscope at 20x magnification and excitation wavelength (488nm) 48h after infiltration. Bar, 50μm. Similar results were obtained in three independent experiments.

### Mutations in the serine-193 and serine-477 containing 14-3-3 protein binding motifs of XopX abolishes the ability of the protein to suppress rice immune responses

In order to understand the functional role of the interaction of XopX with Gf14d and Gf14e, we went ahead to study the role of XopX in the suppression of the rice immune responses. Earlier work in the lab showed that XopX, along with XopQ, XopN and XopZ, is important for the suppression of cell-wall degrading enzyme (CWDE) induced immune responses of rice (Sinha et al., 2013). Since XopX was seen to be binding to the rice 14-3-3 proteins, the effect of mutations in the five 14-3-3 protein binding motifs of XopX on the ability to suppress the rice immune responses was assessed. For this purpose, we took advantage of the observation that a quadruple mutant (QM) strain (Sinha et al., 2013), which is deficient in the production of the XopQ, XopX, XopN and XopZ proteins, induces defence response associated callose deposition and programmed cell death (PCD). Assaying for the PCD indicated that XopX wild-type was able to suppress PCD induced by the QM strain, as signified by internalization of the propidium iodide (PI) stain, whereas mutation to alanine in two of the five 14-3-3 protein binding motifs of *xopX*, motif-2 (*xopX S193A*) and motif-4 (*xopX S477A*), affected the ability of the XopX protein to suppress PCD induced by QM (Fig 3A). However, the phosphomimic mutants *xopX S193D* and *xopX S477D* suppressed PCD induced by the QM (Fig 3A). Similar results were obtained in callose deposition assays in which pHM1∷*xopX,* pHM1*∷xopX S84A,* pHM1*∷xopX S193A,* pHM1*∷xopX S193D,* pHM1*∷xopX T430A*, pHM1∷*xopX S477A,* pHM1*∷xopX S477D* or pHM1*∷xopX T621A* were overexpressed in the QM background. Here, XopX was seen to be able to suppress callose deposition induced by the QM, whereas, XopX S193A as well as XopX S477A were found to be deficient in the suppression of callose deposition induced by the QM (Fig 3B & C). However, both XopX S193D as well as XopX S477D could suppress callose deposition (Fig 3B & C). XopX S84A, XopX T430A and XopX T621A could suppress both callose deposition as well as PCD induced by the QM, as efficiently as wild type XopX, indicating that these motifs are probably not important for suppression of rice immune responses by XopX (Fig 3A-C).

**Fig 3.**
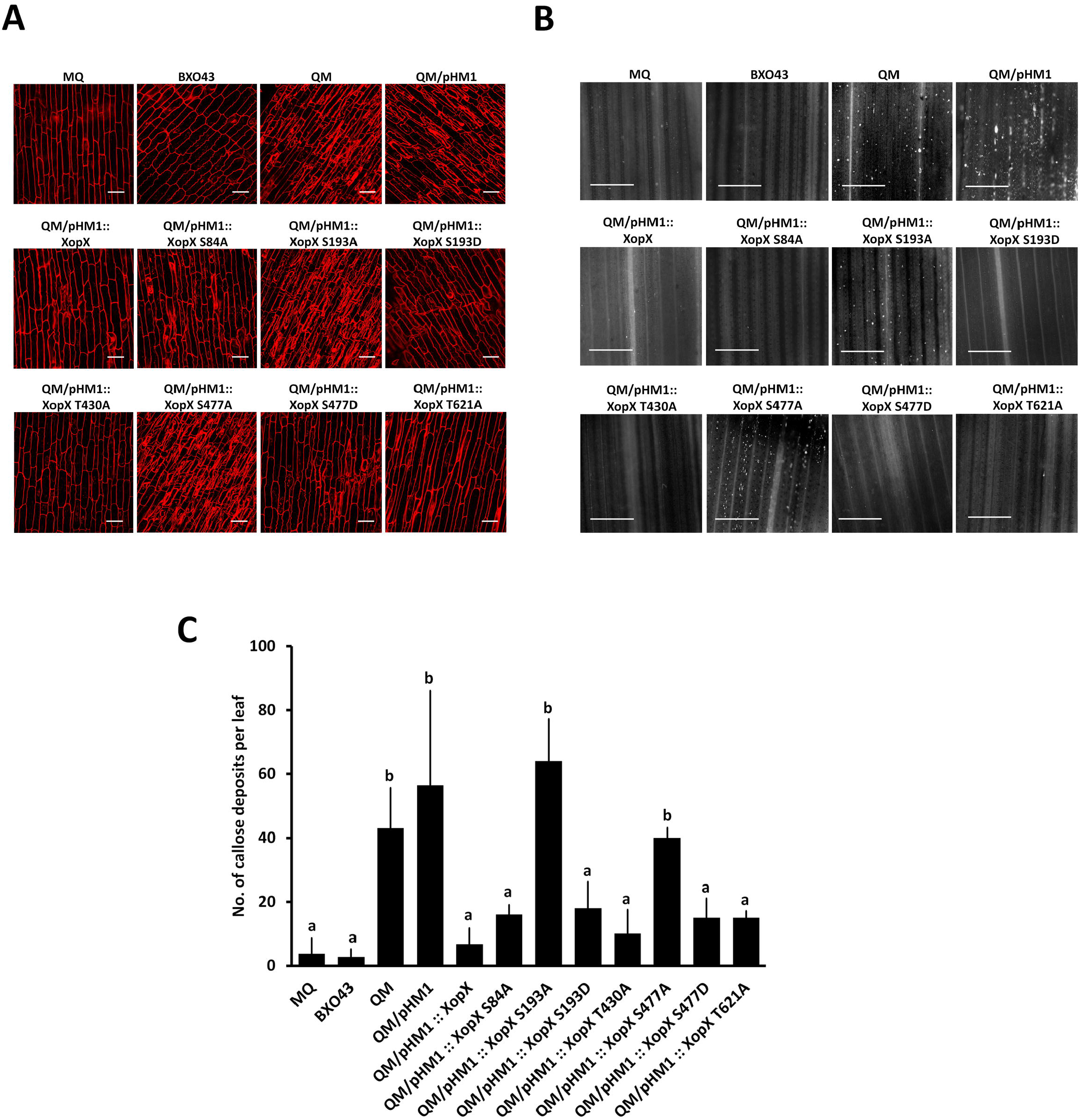
Mutation in the serine-193 and serine-477 containing 14-3-3 protein binding motifs of XopX abolishes its ability to suppress the rice immune responses. **(A)** Rice roots were treated with one of the following: Milli-Q (MQ) water, *X. oryzae* pv. *oryzae* BXO43 (wild-type) or the quadruple mutant (QM) strain harbouring the following: pHM1 empty vector alone, or pHM1 expressing XopX, XopX S84A, XopX S193A, XopX S193D, XopX T430A, XopX S477A, XopX S477D or XopX T621A. Treated roots were subsequently stained with propidium iodide (PI) and observed under a confocal microscope using 63x oil immersion objectives and He-Ne laser at 543nm excitation to detect PI internalization. Five roots were imaged for each construct per experiment. Bar, 20μm. Internalization of PI is indicative of defence response-associated programmed cell death. Similar results were obtained in three independent experiments. **(B & C)** For callose deposition assay, leaves of two-week old rice seedlings were infiltrated with one of the following: MQ water, BXO43, QM strain and QM harbouring the following: pHM1 empty vector alone, or pHM1 expressing XopX, XopX S84A, XopX S193A, XopX S193D, XopX T430A, XopX S477A, XopX S477D or XopX T621A. The leaves were stained 16h later with aniline blue and visualized under an epifluorescence microscope (365nm) at 10x magnification. Mean and standard deviation were calculated for number of callose deposits observed per leaf. Error bars indicate the standard deviation of readings from five infiltrated leaves. Columns in plots capped with the same letter were not significantly different from each other based on analysis of variance done using the Tukey-Kramer honestly significance difference test (*P*< 0.05). Bar, 100μm. Similar results were obtained in three independent experiments.

### Mutation in the serine-193 and serine-477 containing 14-3-3 protein binding motifs of XopX abolishes its ability to localise to the nucleus

Since 14-3-3 proteins are known to alter the subcellular localization of their client proteins, we further checked the subcellular localization of the XopX protein and its 14-3-3 protein binding mutants XopX S193A, XopX S193D, XopX S477A and XopX S477D by tagging them with the eGFP protein and transient overexpression in onion epidermal cells through *A. tumefaciens* AGL1. Visualisation for fluorescence 48h after infection with *A. tumefaciens* expressing the respective eGFP tagged XopX protein or its 14-3-3 protein binding mutants revealed that the XopX wild-type protein localises mostly to the nucleus, but also to the peripheral cytoplasm (Fig 4). However, the 14-3-3 protein binding mutants of XopX which were deficient in 14-3-3 protein binding and immune response suppression (XopX S193A and XopX S477A) were found to be unable to localise to the nucleus, and were seen exclusively in the cytoplasm (Fig 4). The phosphomimic mutants XopX S193D and XopX S477D exhibited a nucleo-cytoplasmic localisation that is similar to wild type XopX (Fig 4).

**Fig 4.**
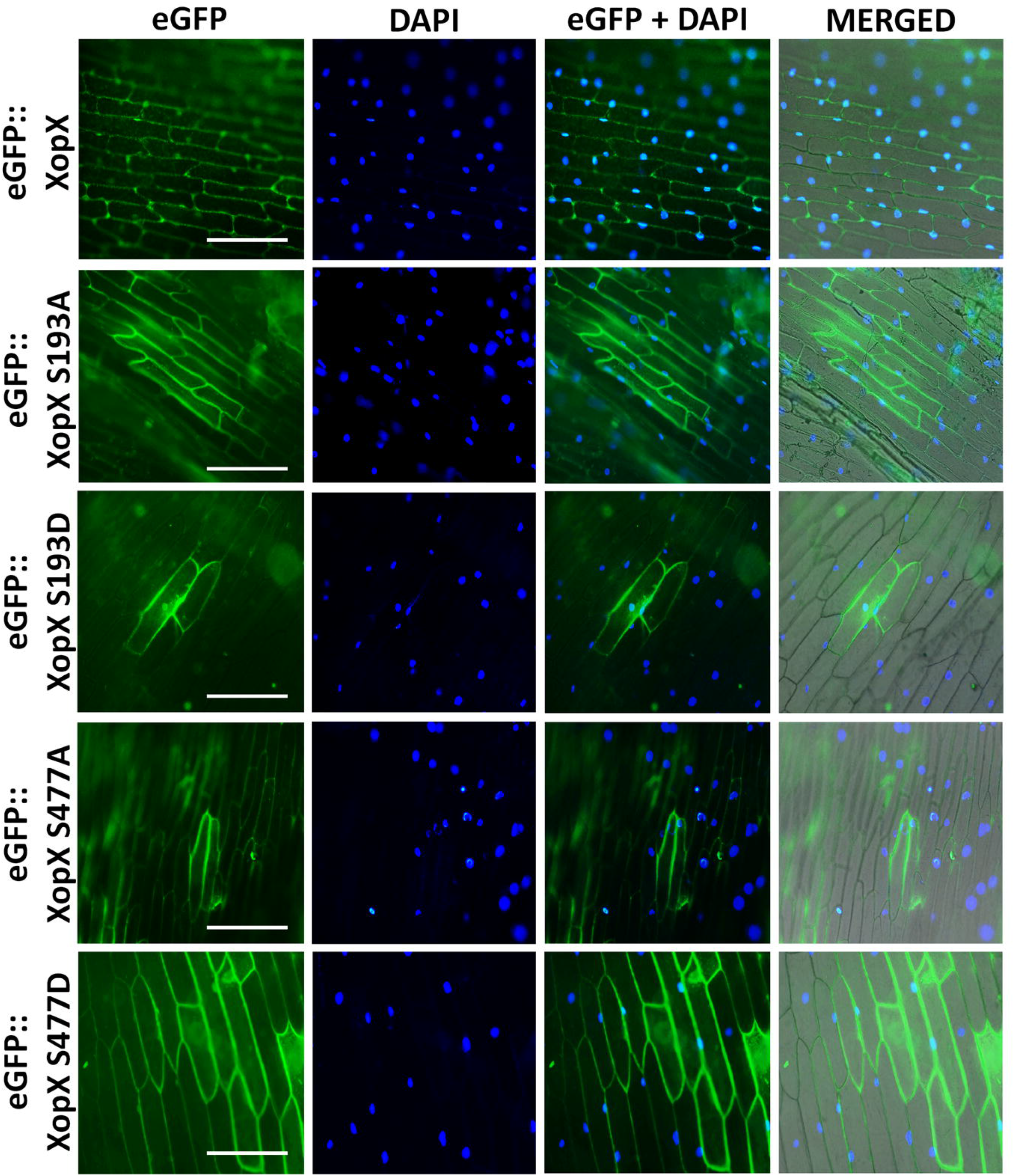
Mutation in the serine-193 and serine-477 containing 14-3-3 protein binding motifs of XopX alters its subcellular localization. *A. tumefaciens* strain AGL1 expressing one of the following was co-cultivated with onion epidermal peels: eGFP∷XopX, eGFP∷XopX S193A, eGFP∷XopX S193D, eGFP∷XopX S477A or eGFP∷XopX S477D. Fluorescence was visualised in an epifluorescence microscope at 10x magnification and excitation wavelength (488nm) 48h after co-cultivation. Bar, 100μm. Similar results were obtained in three independent experiments.

### XopX interacts with XopQ

The XopX protein was found to suppress cell wall degrading enzyme induced immune responses along with the XopN, XopQ and XopZ effector proteins (Sinha et al., 2013). Therefore, we also assessed the ability of the XopX protein to interact with the XopN, XopQ or XopZ proteins using the yeast two-hybrid system. The Activation-domain (AD) fusion clones of *xopQ, xopN* and *xopZ* were made by cloning these genes into the pDEST22 vector yielding *AD∷xopQ*, *AD∷xopN* and *AD∷xopZ* respectively. The *BD∷xopX* clone was used for transformation with *AD∷xopQ*, *AD∷xopN* and *AD∷xopZ* respectively in the yeast strain pJ694a. Primary transformants were selected on synthetic dropout (SD) −leucine −tryptophan double dropout yeast plates. Selection for interaction done on synthetic dropout (SD) - adenine -histidine -leucine -tryptophan quadruple dropout yeast plates supplemented with 1mM 3-AT revealed a positive interaction of XopQ with XopX (Fig 5A). The XopQ-XopX interaction was further confirmed *in-planta* using the BiFC assay, wherein *xopQ* was tagged with the N-terminal portion of the VFP yielding *nVFP∷xopQ* and *xopX* was tagged with the C-terminal portion of VFP yielding *cVFP∷xopX*. Co-cultivation of onion epidermal peels with AGL1 strains expressing these two proteins and further checking for fluorescence revealed strong fluorescence in the nucleus, as seen by co-localisation with DAPI, indicating that the XopX and XopQ proteins interact with each other (Fig 5B). XopQ shows interaction with all the 14-3-3 protein binding motif mutants of XopX, indicating that interaction of XopQ-XopX is not 14-3-3 dependent (Supplementary Fig S1).

**Fig 5.**
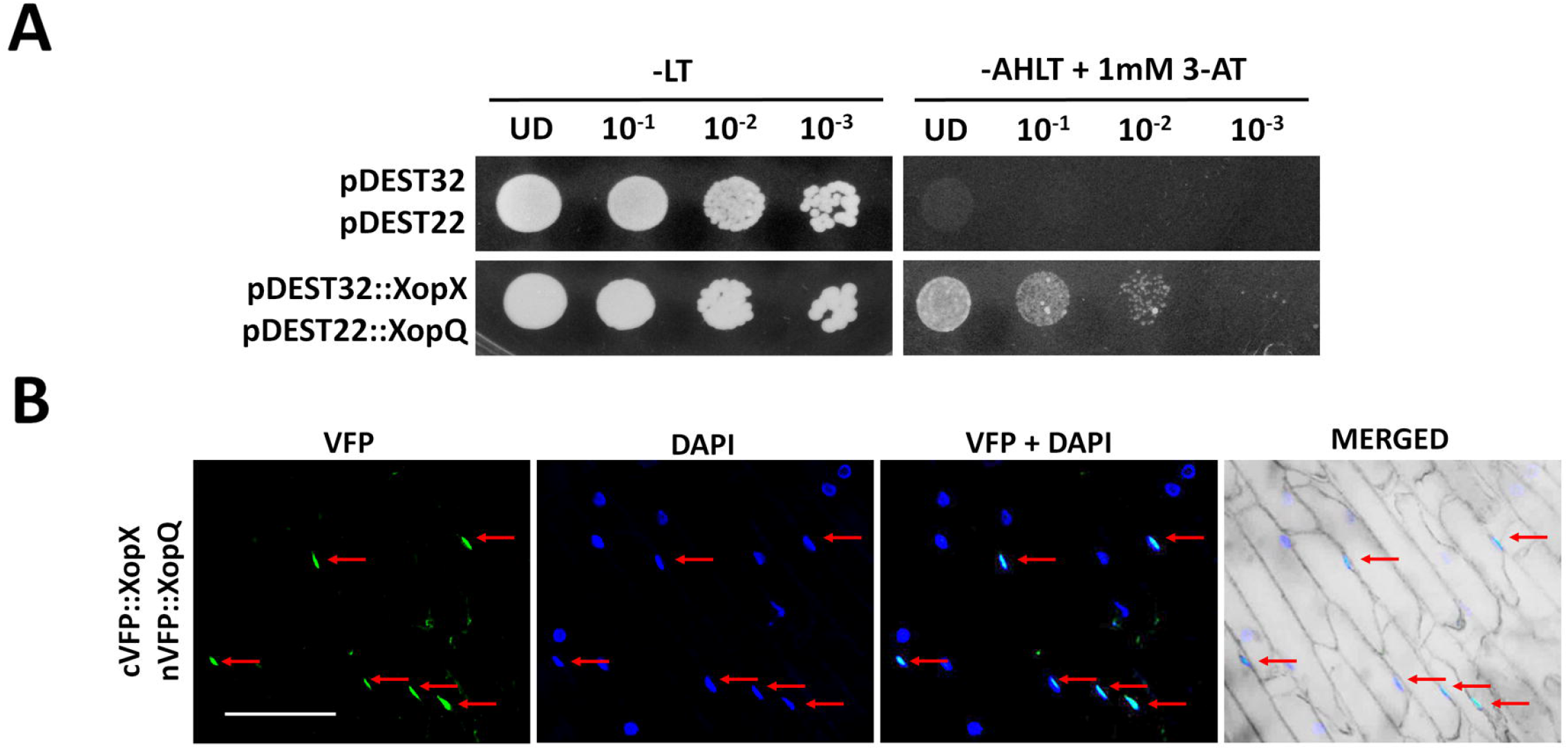
The *X. oryzae* pv. *oryzae* XopQ and XopX proteins interact. **(A)** Yeast strain pJ694a was transformed with pDEST32 vector expressing binding domain (BD) fused with XopX and pDEST22 vector expressing activation domain (AD) fused with XopQ. Transformed colonies were serially diluted and spotted on the nonselective (SD-LT; double dropout DDO) and selective (SD-AHLT; quadruple dropout QDO) media with 1mM 3-AT. Observations were noted after 3 days of incubation at 30°C. Similar results were obtained in three independent experiments. **(B)** For BiFC analysis of XopQ-XopX interactions, onion epidermal peels were co-cultivated with two *A. tumefaciens* AGL1 strains expressing nVFP∷XopQ and cVFP∷XopX. Fluorescence was visualised in an epifluorescence microscope at 10x magnification and excitation wavelength (488nm) 48h after co-cultivation. Bar, 100μm. Similar results were obtained in three independent experiments.

### XopQ-XopX co-expression induces rice immune responses

We transiently overexpressed the eGFP∷XopQ, eGFP∷XopX and eGFP∷XopQ-eGFP∷XopX proteins in rice roots through *A. tumefaciens* AGL1. PI staining for the induction of PCD showed negligible PCD induction after treatment with XopQ and XopX, marked by internalisation of PI (Fig 6A). However, when XopQ and XopX were co-expressed, a high degree of PCD marked by PI internalisation was seen (Fig 6A). This seemed to indicate that co-expression of XopQ and XopX induced PCD in rice roots. A similar result was observed in a callose deposition assay, wherein co-expression of XopQ-XopX, but not of the individual proteins, led to a higher number of callose deposits (Fig 6B & C).

**Fig 6.**
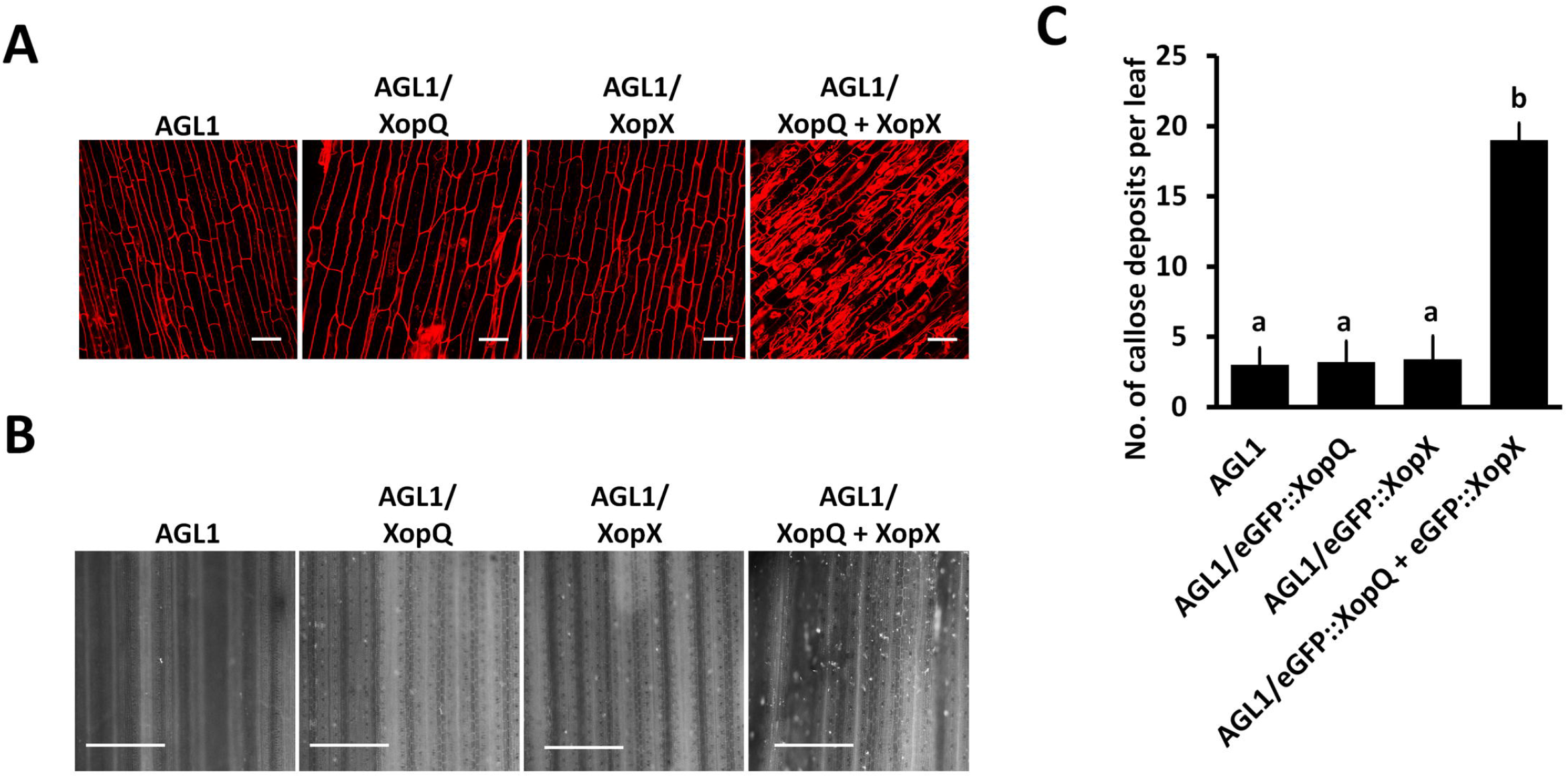
Overexpression of the XopQ-XopX, induces the rice immune responses. **(A)** Rice roots were treated with one of the following: *A. tumefaciens* strain AGL1 alone or AGL1 expressing eGFP∷XopQ, eGFP∷XopX or eGFP∷XopQ + eGFP∷XopX. Treated roots (n=5) were subsequently stained with propidium iodide (PI) and observed under a confocal microscope using a 63x oil immersion objectives and He-Ne laser at 543nm excitation to detect PI internalization. Internalization of PI is indicative of defence response-associated programmed cell death in rice roots. Bar, 20μm. Similar results were obtained in three independent experiments. **(B & C)** For callose deposition assay, leaves of 14-day old rice seedlings were infiltrated with one of the following: *A. tumefaciens* AGL1 alone or AGL1 expressing eGFP∷XopQ, eGFP∷XopX or with a suspension of two strains expressing eGFP∷XopQ + eGFP∷XopX. The leaves were stained 16h later with aniline blue and visualized under an epifluorescence microscope (365nm) at 10x magnification. Mean and standard deviation were calculated for number of callose deposits observed per leaf. Error bars indicate the standard deviation of readings from 5 inoculated leaves. Columns in plots capped with the same letter were not significantly different from each other based on analysis of variance done using the Tukey-Kramer honestly significance difference test (*P*< 0.05). Bar, 100μm. Similar results were obtained in three independent experiments.

### Phosphorylation of XopX at serine-193 and serine-477 is essential for XopQ-XopX induced immune responses

Since the XopX proteins showed interaction with 14-3-3 proteins, we asked if the 14-3-3 proteins would play a role in the induction of immune responses by XopQ-XopX. The effect of mutations of motif-2, eGFP∷XopX S193A, and motif-4, eGFP∷XopX S477A, on XopQ-XopX induced immune responses was assessed. It was observed that overexpression of either the XopX S193A or the XopX S477A mutants, along with XopQ, failed to induce the PCD response in rice (Fig 7A). This is in contrast to induction of immune responses in rice by XopX-XopQ. The phosphomimic mutants XopX S193D and XopX S477D, however, showed induction of PCD along with XopQ (Fig 7A). Similar results were observed in a callose deposition assay (Fig 7B & C). This indicated that phosphorylation was necessary for XopQ-XopX induced immune responses.

**Fig 7.**
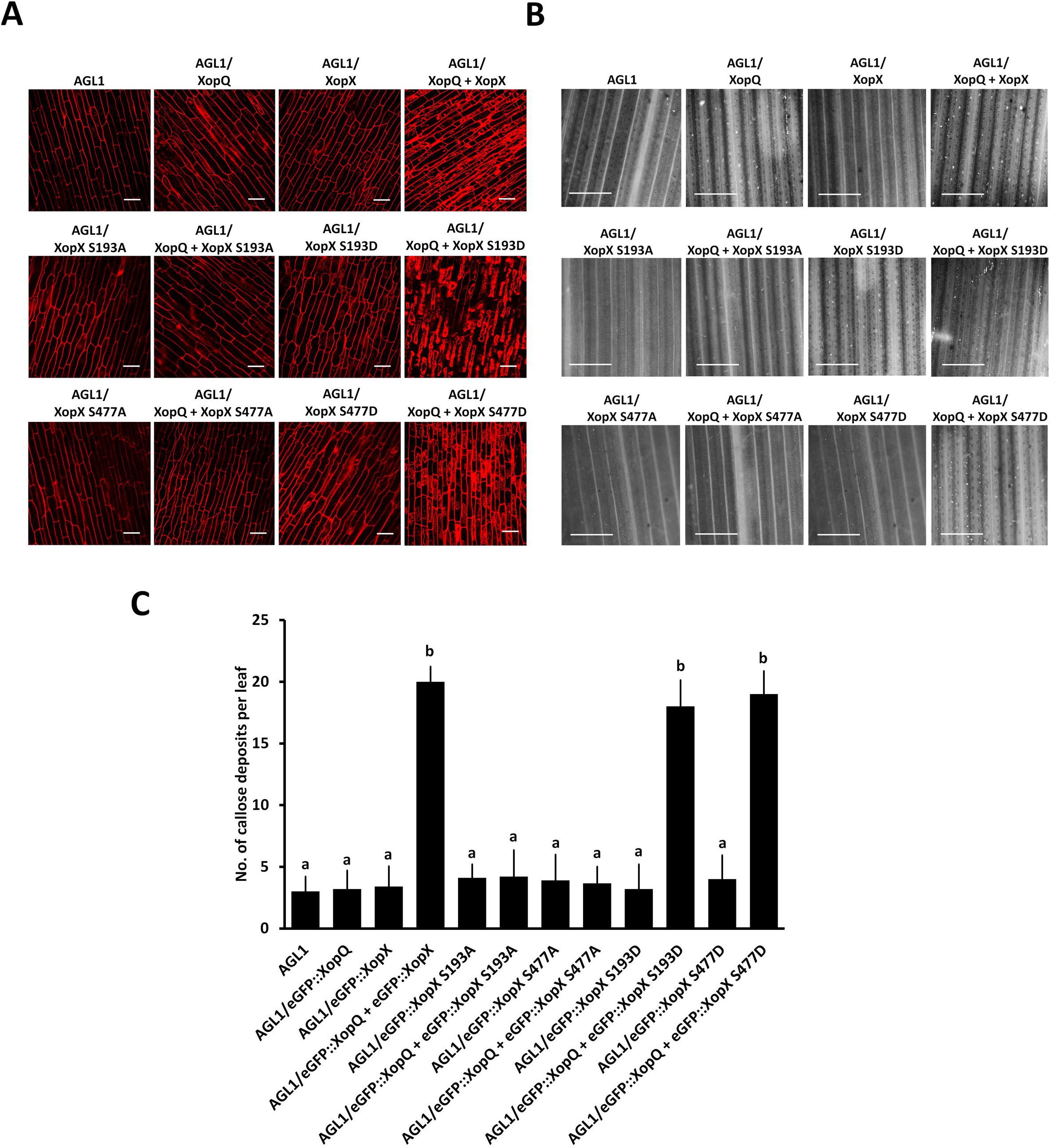
XopX interaction with the rice 14-3-3 proteins is essential for XopQ-XopX induced immune responses. **(A)** Rice roots were treated with *A. tumefaciens* strain AGL1 alone or AGL1 expressing eGFP∷XopQ, eGFP∷XopX, eGFP∷XopQ + eGFP∷XopX, eGFP∷XopX S193A, eGFP∷XopQ + eGFP∷XopX S193A, eGFP∷XopX S193D, eGFP∷XopQ + eGFP∷XopX S193D, eGFP∷XopX S477A, eGFP∷XopQ + eGFP∷XopX S477A, eGFP∷XopX S477D or eGFP∷XopQ + eGFP∷XopX S477D. Treated roots (n=5) were subsequently stained with propidium iodide (PI) and observed under a confocal microscope using a 63x oil immersion objectives and He-Ne laser at 543nm excitation to detect PI internalization. Internalization of PI is indicative of defence response-associated programmed cell death in rice roots. Bar, 20μm. Similar results were obtained in three independent experiments. **(B & C)** For callose deposition assay, leaves of 14-day old rice seedlings were infiltrated with one of the following: *A. tumefaciens* strain AGL1 alone or AGL1 expressing eGFP∷XopQ, eGFP∷XopX, eGFP∷XopQ + eGFP∷XopX, eGFP∷XopX S193A, eGFP∷XopQ + eGFP∷XopX S193A, eGFP∷XopX S193D, eGFP∷XopQ + eGFP∷XopX S193D, eGFP∷XopX S477A, eGFP∷XopQ + eGFP∷XopX S477A, eGFP∷XopX S477D or eGFP∷XopQ + eGFP∷XopX S477D. The leaves were stained 16h later with aniline blue and visualized under an epifluorescence microscope (365nm) at 10x magnification. Bar, 100μm. Mean and standard deviation were calculated for number of callose deposits observed per leaf. Error bars indicate the standard deviation of readings from 5 inoculated leaves. Columns in plots capped with the same letter were not significantly different from each other based on analysis of variance done using the Tukey-Kramer honestly significance difference test (*P*< 0.05). Similar results were obtained in three independent experiments.

### A unique set of type III effectors can suppress XopQ-XopX mediated immune responses

Since the co-expression of XopQ-XopX induced rice immune responses, we asked if any of the other type III effectors could suppress this immune response. For this, 19 type III non-TAL effectors (as listed in Supplementary table S4) were cloned along with the eGFP tag and transiently overexpressed through *A. tumefaciens* AGL1 in rice roots. XopR, XopT and XopAA could not be tested due to difficulty in amplification of the genes. We observed that pre-treatment with 5 of the 19 effectors tested, namely XopU, XopV, XopP, XopG and AvrBs2, could suppress XopQ-XopX induced PCD. We observed significantly lesser internalisation of PI stain after XopQ-XopX treatment when rice roots were pre-treated with these five type III effectors, as opposed to XopQ-XopX treatment alone (Fig 8A). This indicated that these effectors could successfully suppress the PCD induced by XopQ-XopX treatment. Overexpression of XopG, XopP, XopU, XopV or AvrBs2 individually did not induce the PCD response in rice (Fig 8A). Expression of these effectors was also checked in *N. tabacum* (Supplementary Fig S2). Pre-treatment with the other 14 effectors, XopI, XopK, XopL, XopAD, HpaA, XopW, XopA, XopL, XopAE, XopC, XopF, XopAB, XopN, XopZ or XopY were seen to be unable to suppress the PCD induced by overexpression of XopQ-XopX (Supplementary Fig S3). All 14 of these effectors were found to be expressing, as visualised in *N. tabacum* (Supplementary Fig S4). This was further confirmed using a callose deposition assay, wherein similar results were obtained. XopG, XopP, XopU, XopV and AvrBs2 were found to be able to suppress the callose deposition induced by treatment with XopQ and XopX (Fig 8 B & C).

**Fig 8.**
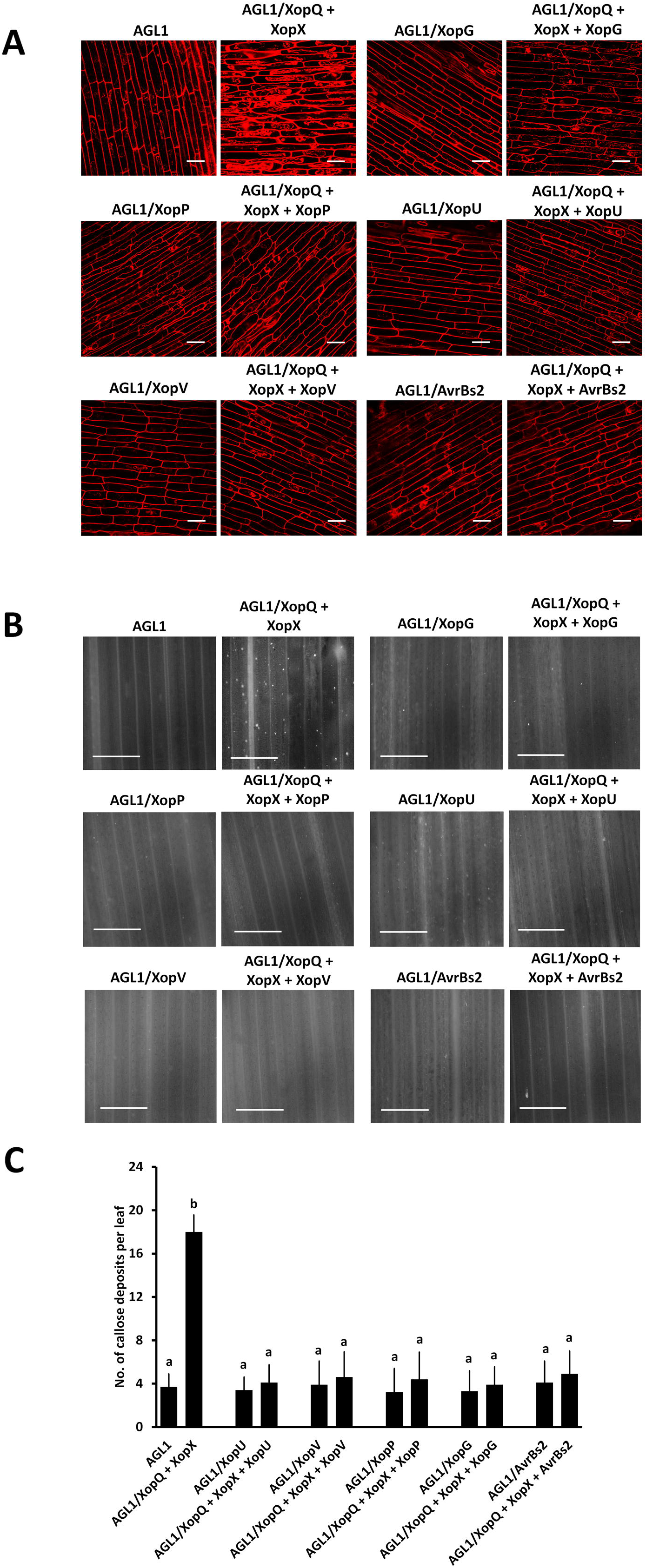
XopG, XopP, XopU, XopV and AvrBs2 can suppress XopQ-XopX induced immune responses. **(A)** Rice roots were treated with *A. tumefaciens* strain AGL1 alone or AGL1 expressing *A. tumefaciens* AGL1 alone or AGL1 expressing eGFP∷XopQ + eGFP∷XopX, eGFP∷XopG, eGFP∷XopP, eGFP∷XopU, eGFP∷XopV or eGFP∷AvrBs2, or pre-treatment with eGFP∷XopG, eGFP∷XopP, eGFP∷XopU, eGFP∷XopV or eGFP∷AvrBs2, followed by overexpression of eGFP∷XopQ + eGFP∷XopX. Treated roots (n=5) were subsequently stained with propidium iodide (PI) and observed under a confocal microscope using a 63x oil immersion objectives and He-Ne laser at 543nm excitation to detect PI internalization. Internalization of PI is indicative of defence response-associated programmed cell death in rice roots. Bar, 20μm. Similar results were obtained in three independent experiments. **(B & C)** For callose deposition assay, leaves of 14-day old rice seedlings were infiltrated with one of the following: *A. tumefaciens* AGL1 alone or AGL1 expressing eGFP∷XopQ + eGFP∷XopX, eGFP∷XopG, eGFP∷XopQ + eGFP∷XopX + eGFP∷XopG, eGFP∷XopP, eGFP∷XopQ + eGFP∷XopX + eGFP∷XopP, eGFP∷XopU, eGFP∷XopQ + eGFP∷XopX + eGFP∷XopU, eGFP∷XopV, eGFP∷XopQ + eGFP∷XopX + eGFP∷XopV, eGFP∷AvrBs2 or eGFP∷XopQ + eGFP∷XopX + eGFP∷AvrBs2. The leaves were stained 16h later with aniline blue and visualized under an epifluorescence microscope (365nm) at 10x magnification. Bar, 100μm. Mean and standard deviation were calculated for number of callose deposits observed per leaf. Error bars indicate the standard deviation of readings from 5 inoculated leaves. Columns in plots capped with the same letter were not significantly different from each other based on analysis of variance done using the Tukey-Kramer honestly significance difference test (*P*< 0.05). Similar results were obtained in three independent experiments.

## Discussion

This study highlights the dual role of the XopQ-XopX type III effectors in modulation of immune responses of rice. The *X. oryzae* pv. *oryzae* XopX has earlier been shown to be important for the suppression of cell-wall degrading enzyme induced immune responses in rice (Sinha et al., 2013). XopX from *X. campestris* pv. *vesicatoria* also contributes to the virulence of the bacteria on the host plants, pepper and tomato, and modulates the plant immune response, resulting in enhanced susceptibility of the plant (Metz et al., 2005). However, the XopX effector protein from *Xanthomonas campestris* pv. *vesicatoria (Xcv)* has also been shown to be an inducer of rice immune responses wherein it was shown to weakly upregulate PTI marker genes (Stork et al., 2015). In the current work, we demonstrate that XopX suppresses rice immune responses, by interaction with the 14-3-3 proteins. XopX interacts with two rice 14-3-3 proteins, namely Gf14d And Gf14e. Both motif-2 encompassing XopX S193 as well as motif-4 encompassing XopX S477 seem to be required for the interaction of XopX with the 14-3-3 proteins as XopX loses interaction with both Gf14d and Gf14e if either motif-2 or motif-4 is mutated to a phospho-null motif. Hence, it appears that both motifs are required for interaction with Gf14d and Gf14e, which had earlier been shown to be a negative regulator of rice immune responses. The role of Gf14d in elaboration of rice immune responses is not yet established. The observation that XopX mutations that are unable to interact with Gf14d and Gf14e are also unable to suppress rice immune responses suggests that at least one of these two 14-3-3 proteins might be a positive regulator of rice immune responses. As Gf14e is shown to be a negative regulator of rice immune responses, it is possible that Gf14d is a positive regulator of rice immune innate immunity. However, this needs to be established. The significance of the interaction of XopX with a negative regulator of innate immunity such as Gf14e is also to be investigated. Another question which arises is also that, since both XopQ and XopX are suppressors of immune responses, is there a temporal or spatial distribution of XopQ and XopX which decides the interaction and hence suppression/ induction of immune responses? XopQ is a highly conserved core effector in both *X. oryzae* pv. *oryzae* as well as *X. oryzae* pv. *oryzicola* strains (Midha et al., 2017, Hajri et al., 2009). XopQ and XopX both have a nucleo-cytoplasmic localisation. It has been shown earlier that the mutants of XopQ which were deficient in 14-3-3 protein binding (XopQ S65A), fails to localise to the cytoplasm. On the other hand, mutants of XopX which were deficient in 14-3-3 protein binding (XopX S193A and XopX S477A) fail to localise to the nucleus. Hence, phosphorylation, and interaction with the cognate 14-3-3 proteins seem to be important for cytoplasmic localisation of XopQ and nuclear localisation of XopX and this localization might be important for their activity in suppression of innate immunity. This leads us to hypothesise that this nucleo-cytoplasmic partitioning may be crucial for suppression of immune responses. The function of suppression of immune responses might be specifically taking place in the cytoplasm in case of XopQ and in the nucleus for XopX. However, interaction of XopQ and XopX may sequester these two proteins, leading to loss of suppression of immune responses. Their interaction and co-localisation may also be activating downstream pathways for induction of immune responses (Supplementary Fig S5). There may also be a temporal distribution in expression of XopQ and XopX, as XopX has been shown to be expressed in early stages of host infection at 3 and 6 days after infection (Soto-Suarez et al., 2010). Such dual behaviour of T3SS effectors in supressing and inducing plant immune responses has also been reported previously for the *Pseudomonas syringae* pv. *tomato (Pst)* effector AvrE1 and for the *Xanthomonas euvesicatoria* XopX (Badel et al., 2006, Stork et al., 2015).

It is a possibility that the XopQ-XopX induced immune responses is a form of defence response that is triggered by these two effectors. The interaction of XopQ and XopX may be sensed by the rice plant, which then might be able to mount an effector triggered immune response. In such a scenario, it may be imperative for *X. oryzae* pv. *oryzae* to be able to suppress this effector triggered defence response. Indeed, we find that five effectors, XopU, XopV, XopP, XopG and AvrBs2, out of the nineteen non-TAL effectors screened, could suppress XopQ-XopX induced immune responses. Out of these five effectors, AvrBs2 and XopV are part of the core effectors in 113 strains of Indian, Asian, African and USA strains of *X. oryzae* (Midha et al., 2017), and are also common between *X. oryzae* pv. *oryzae* and the bean pathogen, *X. axonopodis* pv. *phaseoli* (Aritua et al., 2015). XopP, XopU and XopV have also been shown to be ubiquitously present in *X. oryzae* pv. *oryzae* and *X. oryzae* pv. *oryzicola* strains (Hajri et al., 2009). AvrBs2 from *X. oryzae* pv. *oryzicola* has earlier been shown to be an essential virulence factor that contributes to bacterial virulence and multiplication by inhibiting the rice defence responses (Li et al., 2015) and is highly conserved among strains of *X. campestris* pv. *vesicatoria*, and other *X. campestris* pathovars (Kearney and Staskawicz, 1990). The *X. oryzae* pv. *oryzae* XopP protein has been shown to interacts with the U-box domain of an E3 ubiquitin ligase OsPUB44, thereby inhibiting the E3 ubiquitin ligase activity of OsPUB44 (Ishikawa et al., 2014). Interestingly, preliminary bioinformatic analysis has revealed that XopU, XopV, XopP, XopG and AvrBs2, all have at least one putative mode-I 14-3-3 protein binding motif with a conserved serine [XopU: ‘RAESTP’, amino acid 374-379; XopV: ‘RIRSTP’, amino acid 142-147; XopP: ‘RLESLP’, amino acid 494-499; XopG: ‘RLGSNP’, amino acid 70-75; AvrBs2: ‘RAVSIP’, amino acid 46-51 and ‘RAASGP’, amino acid 143-148]. Hence, it is possible that these effectors may suppress XopQ-XopX induced immune responses by interaction with the rice 14-3-3 proteins.

Thus, this study suggests a dual role for the type III effectors XopQ and XopX in the course of disease progression of *X. oryzae* pv. *oryzae* both as suppressor and inducers of immune responses in rice, and identifies bacterial effectors that may be involved in suppression of effector-triggered immunity.

Is there a temporal distribution in expression of XopQ and XopX during progression of disease caused by *X. oryzae* pv. *oryzae*? What are the roles of the 14-3-3 proteins Gf14d, Gf14e, Gf14f and Gf14g in elaboration of rice immune responses during *X. oryzae* pv. *oryzae* infection? What are the additional rice factors which might regulate the localisation of these proteins and their interaction? These questions, and elucidation of the mechanism and biological significance of the interaction XopQ and XopX are aspects which require further research.

## Experimental Procedures

### Bacterial strains and plant material

The bacterial strains *Escherichia coli* DH5□; *Agrobacterium tumefaciens* AGL1, *X. oryzae* pv. *oryzae* strain BXO43 (Thieme et al., 2005) and the *X. oryzae* pv. *oryzae* mutant ∆*xopQ xopN-* ∆*xopX* ∆*xopZ* quadruple mutant (QM) (Sinha et al., 2013) were used for the study. *E. coli* and *A. tumefaciens* AGL1 strain were grown in Luria–Bertani (LB) medium. *E. coli* was grown at 37°C whereas *A. tumefaciens* was grown at 28°C. *X. oryzae* pv. *oryzae* strains were grown on peptone sucrose (PS) medium at 28°C (Ray et al., 2000). The yeast strain pJ694a was grown at 30°C in yeast extract, peptone, dextrose (YPD) medium, or minimal media supplemented with suitable amino acids for auxotrophic selection. The plant cultivars used were the susceptible rice variety Taichung Native-1 (TN-1) for transient overexpression studies in rice and *Nicotiana benthamiana* or *Nicotiana tabacum* for ectopic overexpression of proteins for bimolecular fluorescence complementation assay or expression analysis. The concentrations of antibiotics used were rifampicin (Rif)-50μg/ml, spectinomycin (Sp)-50μg/ml, gentamycin (Gent)-10μg/ml, ampicillin (Amp)-100μg/ml, kanamycin (Km)-15μg/ml for *X. oryzae* pv. *oryzae* and 50μg/ml for *E. coli*.

### Rice growth conditions

The TN-1 rice variety, which is susceptible to *X. oryzae* pv. *oryzae* infection, was used to study bacterial leaf blight symptoms caused by the *X. oryzae* pv. *oryzae* strain BXO43. Healthy seeds of the plants were surface-sterilized in sodium hypochlorite (Sigma) for 2 min, rinsed five times in deionized water and imbibed overnight at 28°C. They were then placed on moist filter paper for 2 days in dark at 28°C. Upon emergence of root and shoot, they were transferred to 14hr light/ 10hr dark photoperiod in growth chamber (Conviron, Germany). After 3 days of growth, seedlings were sown in black soil mix. Pots with plants were kept in a greenhouse in the following conditions: ~30°C/20°C (day/night), ~80% humidity, natural sunlight with a ~13h/11h light/dark photoperiod.

### Molecular biology and microbiology techniques

For the amplification and cloning of the wild-type copy of the *xopX* gene, or its 14-3-3 protein binding motif mutants, high-fidelity Phusion polymerase (Finnzymes) was used along with their respective primers (Supplementary table S1). The genes were cloned into pENTR/D-TOPO (Invitrogen, California) and further by Gateway LR reaction (Invitrogen, California) into Gateway compatible vectors. Taq polymerase ReadyMix (KAPA Biosystems, Wilmington, MA) was used for all screening purposes. For cloning in the pHM1 vector, primers as listed in Supplementary table S1 were used for amplification of the *xopX* gene and its 14-3-3 protein binding motif mutants using Phusion polymerase (Thermo Fischer Scientific, Massachusetts). Restriction digestions were carried out using Fast Digest enzymes (Thermo Fischer Scientific, Massachusetts) specific to the restriction enzyme sites included in the primers. Ligation reactions for cloning in pHM1 were carried out using T4 DNA ligase (NEB, Massachusetts).

Plasmids were purified using the alkaline lysis method. Gel extractions were carried out using Macherey Nagel Gel Extraction kits. Agarose gel electrophoresis, transformation of *E. coli* and electroporation of plasmids into *A. tumefaciens* AGL1 and *X. oryzae* pv. *oryzae* were performed as described previously (Ray et al., 2000, Subramoni and Sonti, 2005). All cloned vectors (Supplementary table S2) were confirmed by sequencing (ABI Prism 3700 automated DNA sequencer). The obtained sequences were subjected to homology searches using the BLAST algorithm in the National Centre for Biotechnology Information database (Altschul et al., 1990). Site-directed mutagenesis was done in *xopX* based on prediction of the 14-3-3 protein binding motifs in *xopX.* The conserved serine/threonine residues in the interaction motifs were mutagenized to alanine to yield a null mutant and to aspartic acid to yield a phosphomimic mutant using primers in Supplementary table S1. The pENTR∷*xopX* plasmid was used as template.

### Yeast two-hybrid assays

The wild-type copy of the *xopX* gene and its 14-3-3 protein binding motif mutants were cloned in the yeast two-hybrid vector pDEST32 (Invitrogen) using the Gateway cloning system (Invitrogen, California). The eight rice *14-3-3* genes cloned in the yeast two-hybrid vector pDEST22 (Invitrogen) were used from a previous study (Deb et al., 2019). For analysis of interaction of *xopX* with the other effector proteins, the pDEST22 clones containing *xopN*, *xopQ* and *xopZ* were used, whereas *xopX* was cloned in pDEST32. These plasmids were transformed into *Saccharomyces cerevisiae* strain pJ694a (James et al., 1996). Yeast transformation was done using the LiAc/single strand carrier DNA/PEG method as described by Geitz *et.* al (Gietz and Schiestl, 2007) with changes. Briefly, yeast cells were grown overnight in YPAD medium (1% (w/v) Bacto-yeast extract, 2% (w/v) Bacto-peptone, adenine hemisulfate (80 mg/l). Following 12-16h of growth, secondary culture was put using 3% of primary inoculum and the culture was allowed to grow for 4-6h till it reached to O.D._600_= 0.6-0.8. Cells were then harvested by centrifugation at 3000rpm for five minutes, washed with sterile water and resuspended in sterile water. 360μl of transformation mix (40% PEG3350, 100mM LiAc/TE, 20μg single-stranded carrier DNA and 1μg of each plasmid DNA) was added per plasmid to be transformed, mixed by vortexing vigorously, and incubated at 30°C for 30 minutes. The cells were then subjected to heat shock at 42°C for 30 minutes, placed on ice for 5 minutes and plated on selection medium for the respective transformed vector and grown at 30°C to select for transformants.

For screening for interaction, colonies were scraped from plates, patched and grown overnight in liquid medium with selection at 30°C with shaking. The OD_600_ of saturated cultures was adjusted to 1.0, serial dilutions were made and spotted on the medium with selection for vector and the medium with selection for interaction (lacking the products of the reporter genes adenine and histidine) + 1mM 3-amino triazole (3-AT; inhibitor of His3 gene). Growth on the medium for interaction was used to identify the interacting clones. Each set was repeated three times.

### Bimolecular fluorescence complementation (BiFC)

The wild-type copy of *xopX* and its 14-3-3 protein binding motif mutants were cloned by Gateway cloning (Invitrogen, California) from the pENTR clones to the BiFC vector pDEST-VYCE(R)GW carrying the C-terminal region of the Venus Fluorescent Protein (VFP) (Gehl et al., 2009) by Gateway® cloning (Invitrogen, California) from the pENTR clones to yield the constructs as listed in Supplementary table S2. The eight rice *14-3-3* genes cloned in the BiFC vector pDEST-VYNE(R)GW carrying the N-terminal region of VFP were used from a previous study (Deb et al., 2019). To check for XopQ-XopX interaction, *xopQ* was cloned in pDEST-VYNE(R)GW. These binary vectors obtained were then electroporated into the *A. tumefaciens* strain AGL1 (Supplementary table S2). A suspension of two strains expressing the gene-nVFP/cVFP fusions were grown to 0.8 O.D._600_, resuspended in infiltration buffer (10mM MES, 10mM MgCl_2_, 100μM acetosyringone, pH 5.6) and used for transient expression in *N. benthamiana*. VFP signals were examined 48h after infiltration under a LSM880 confocal microscope (Carl Zeiss, Germany) using 20x objectives and He-Ne laser at 488nm excitation. Images were analyzed using the ZEN software. Each set was repeated three times.

### Callose deposition in rice

Callose deposition assays were done as described earlier (Adam and Somerville, 1996, Hauck et al., 2003, Sinha et al., 2013). *X. oryzae* pv. *oryzae* strains were grown to saturation, OD_600_ adjusted to 1.0 using Milli-Q water and infiltrated with a needleless 1ml syringe into leaves of 14-day old rice plants. 16h after infiltration, the leaves were cut, spanning 0.5 cm on each side of the infiltration zone, and placed in absolute alcohol at 65°C to remove chlorophyll completely. This was followed by treatment with 70% ethanol at 65°C and further by MQ water for rehydration. Subsequently, the samples were stained with 0.05% aniline blue solution prepared in 150mM K_2_HPO_4_, pH 9.5. The leaves were then washed with MQ water and observed under an epifluorescence microscope (Nikon, Japan) using a blue filter (excitation wavelength of 365 nm) and 10x objective. Number of callose were counted per leaf, excluding the zone of infiltration. At least five such leaves were imaged for each construct per experiment. Each set was repeated three times.

### Defence response associated programmed cell death assay

Assays for programmed cell death in rice roots were performed as described earlier (Sinha et al., 2013). TN-1 rice seeds were surface sterilised by washing with sodium hypochlorite (Sigma) followed by three water washes and imbibed with water overnight. The following day, the seeds were placed for germination on 0.5% sterile agar lined with sterile Whatman filter paper for 2 days in dark at 28°C. 1cm long root tips were cut from the seedlings and treated with either *X. oryzae* pv. *oryzae* (strains grown to saturation and O.D._600_ adjusted to 1.0 using MQ water) or *A. tumefaciens* AGL1 harbouring the *eGFP∷gene* fusions (strains grown to saturation and O.D._600_ adjusted to 0.8 using infiltration buffer: 10mM MES, 10mM MgCl_2_, 100μM acetosyringone, pH 5.6). After incubation for 16h, roots were washed and stained with propidium iodide (PI). The samples were visualised under a LSM-880 confocal microscope (Carl Zeiss, Germany) using 63x oil immersion objectives and He-Ne laser at 543nm excitation to detect PI internalization. Images were analyzed using the LSM software. At least five roots were imaged for each construct per experiment. Each set was repeated three times.

### Transient protein expression for localisation in onion epidermal peels

Healthy onion scales (1×1 cm) were placed on plate in such a way that their inner surfaces were immersed in *A. tumefaciens* AGL1 containing the respective *eGFP-gene* fusions (O.D._600_ = 1–1.5) resuspended in a solution consisting of 5% (w/v) sucrose, 100 mg acetosyringone/L and 0.02% (v/v) Silwet-77 for 12h at 28°C. After 12h of incubation, the onion scales were transferred to plates of 1/2 MS (Murashige and Skoog salts, 30 g sucrose/L and 0.7% (g/v) agar, pH 5.7) and co-cultivated with *A. tumefaciens* for 2 days. For visualisation of fluorescence, epidermal peels of the onion scales were carefully removed using a pair of forceps, stained with DAPI stain (2μg/ml) and mounted on slide using 40% glycerol. Fluorescence was visualised under an epifluorescence microscope (Nikon, Japan) at 488nm excitation and 10x objective. Each set was repeated three times.

### Transient protein expression in *Nicotiana tabacum*

The type III effectors were cloned by Gateway cloning (Invitrogen, California) from the pENTR clones to the pH7WGF2 vector containing the eGFP gene (Karimi et al., 2002) by Gateway® cloning (Invitrogen, California) from the pENTR clones to yield the constructs as listed in Supplementary table S2. These binary vectors obtained were then electroporated into the *A. tumefaciens* strain AGL1 (Supplementary table S2). The strain expressing the eGFP∷gene fusions were grown to O.D._600_ = 0.8, resuspended in infiltration buffer (10mM MES, 10mM MgCl_2_, 100μM acetosyringone, pH 5.6) and used for transient expression in *N. tabacum*. eGFP signals were examined 48h after infiltration under a LSM880 confocal microscope (Carl Zeiss, Germany) using 20x objectives and He-Ne laser at 488nm excitation. Images were analyzed using the ZEN software. Each set was repeated three times.

## Acknowledgements

This work was supported by grants to RVS from the Plant-Microbe and Soil Interaction (PMSI) project of the Council of Scientific and Industrial Research (CSIR), Government of India and the J. C. Bose fellowship to RVS from the Department of Science and Technology (DST), Government of India. This work was also supported by grants to HKP from the Council of Scientific and Industrial Research (CSIR), Government of India. SD acknowledges the Council of Scientific and Industrial Research (CSIR), Government of India for Ph.D. fellowship. PG acknowledges the Council of Scientific and Industrial Research (CSIR), Government of India for fellowship.

## Data availability statement

The authors declare that all data has been included in the manuscript.

## Short legends for supporting information

**Supplementary Fig S1. The 14-3-3 protein binding motif mutants of XopQ and XopX interact.** Yeast strain pJ694a was transformed with pDEST32 vector expressing binding domain (BD) fused with XopX, XopX S84A, XopX S193A, XopX S193D, XopX T430A, XopX S477A, XopX S477D or XopX T621A, and pDEST22 vector expressing activation domain (AD) fused with XopQ. Transformed colonies were serially diluted and spotted on the nonselective (SD-LT; double dropout DDO) and selective (SD-AHLT; quadruple dropout QDO) media with 1mM 3-AT. Observations were noted after 3 days of incubation at 30°C. Similar results were obtained in three independent experiments.

**Supplementary Fig S2. Expression of effectors which can suppress XopQ-XopX induced immune responses.** Leaves of *N. tabacum* were syringe-infiltrated with a suspension of *A. tumefaciens* AGL1 strain expressing eGFP∷AvrBs2, eGFP∷XopV, eGFP∷XopG, eGFP∷XopP or eGFP∷XopU. Fluorescence was visualised in a confocal microscope at 20x magnification and excitation wavelength (488nm) 48h after infiltration. Bar, 50μm. Similar results were obtained in three independent experiments.

**Supplementary Fig S3. A subset of type III effectors is unable to suppress XopQ-XopX induced PCD.** Rice roots were treated with *A. tumefaciens* strain AGL1 alone or AGL1 expressing eGFP∷XopQ + eGFP∷XopX, eGFP∷XopN, eGFP∷XopZ, eGFP∷XopY, eGFP∷XopA, eGFP∷XopAB, eGFP∷XopAE, eGFP∷XopC, eGFP∷XopI, eGFP∷XopK, eGFP∷XopL, eGFP∷XopW, eGFP∷HpaA, eGFP∷XopAD or eGFP∷XopF, or pre-treatment with eGFP∷XopN, eGFP∷XopZ, eGFP∷XopY, eGFP∷XopA, eGFP∷XopAB, eGFP∷XopAE, eGFP∷XopC, eGFP∷XopI, eGFP∷XopK, eGFP∷XopL, eGFP∷XopW, eGFP∷HpaA, eGFP∷XopAD or eGFP∷XopF, followed by overexpression of eGFP∷XopQ + eGFP∷XopX. Treated roots (n=5) were subsequently stained with propidium iodide (PI) and observed under a confocal microscope using a 63x oil immersion objectives and He-Ne laser at 543nm excitation to detect PI internalization. Internalization of PI is indicative of defence response-associated programmed cell death in rice roots. Bar, 20μm. Similar results were obtained in three independent experiments.

**Supplementary Fig S4. Expression of effectors which are unable to suppress XopQ-XopX induced immune responses.** Leaves of *N. tabacum* were syringe-infiltrated with a suspension of *A. tumefaciens* AGL1 strain expressing eGFP∷XopI, eGFP∷XopK, eGFP∷XopL, eGFP∷XopAD, eGFP∷HpaA, eGFP∷XopW, eGFP∷XopA, eGFP∷XopAE, eGFP∷XopC, eGFP∷XopF, eGFP∷XopAB, eGFP∷XopN, eGFP∷XopZ or eGFP∷XopY. Fluorescence was visualised in a confocal microscope at 20x magnification and excitation wavelength (488nm) 48h after infiltration. Bar, 50μm. Similar results were obtained in three independent experiments.

**Supplementary Fig S5. Model explaining induction of immune responses by XopQ-XopX.** Cell wall damage is perceived by the host plant to induce a cascade of cellular responses which finally lead to the activation of immune responses in rice. The 14-3-3 proteins are putative activators of the rice immune responses. XopQ and XopX interact with their cognate 14-3-3 partners Gf14f/g or Gf14d/e respectively to suppress the plant immune responses. However, interaction of XopQ and XopX leads to the activation of the rice immune responses, which can be further suppressed by XopG, XopP, XopU, XopV and AvrBs2. The molecules marked in red are the putative candidates involved in the activation of rice immune responses.

**Supplementary table S1.** List of primers used in the study

**Supplementary table S2.** List of plasmids used in the study

**Supplementary table S3.** List of strains used in the study

**Supplementary table S4.** Details of annotated *X. oryzae* pv. *oryzae* effectors screened for suppression of XopQ-XopX mediated immune responses

