## Supplementary figures and images for "Interaction of the *Xanthomonas* effectors XopQ and XopX results in induction of rice immune responses"

### Supplementary Fig S1

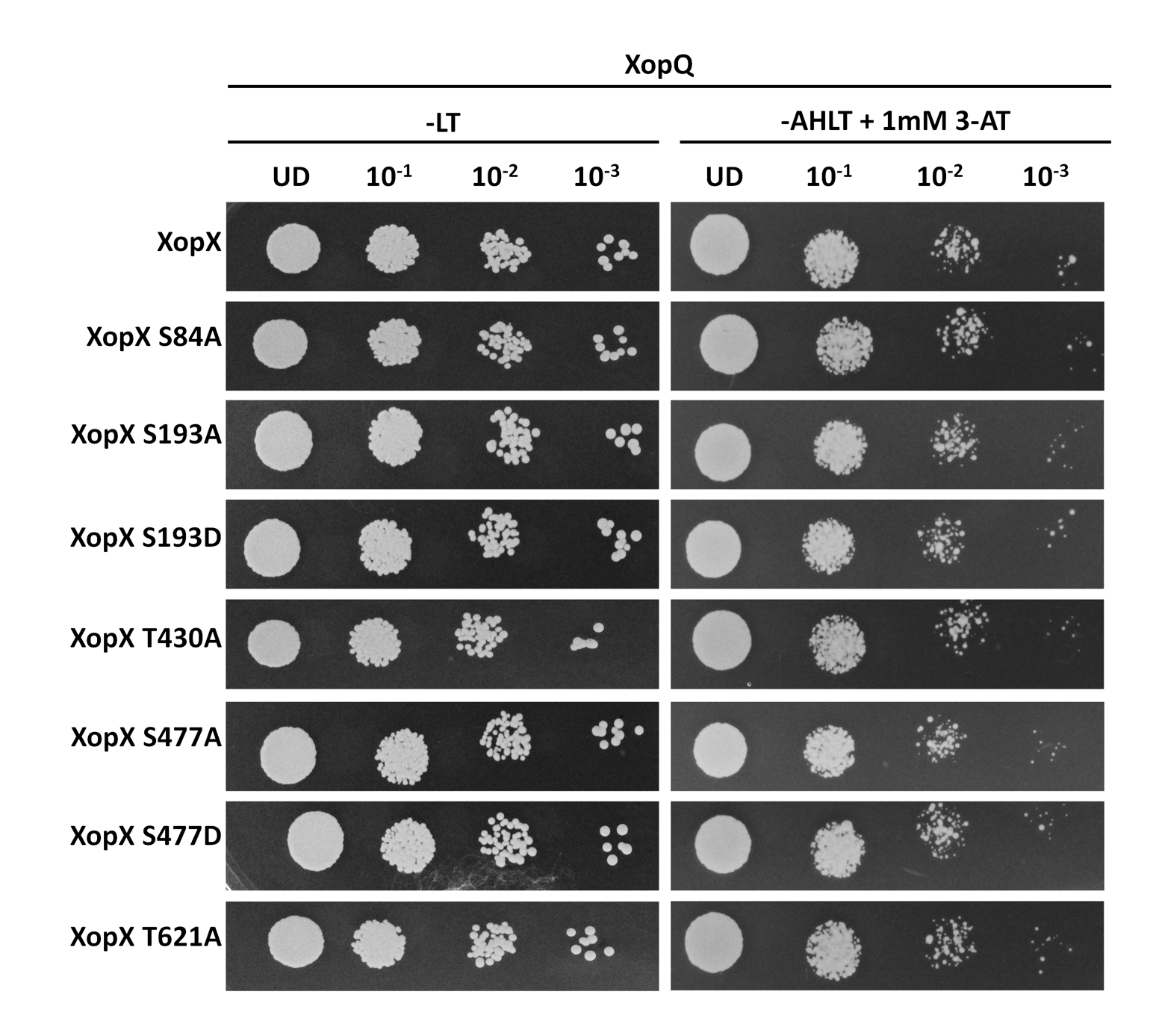

### Supplementary Fig S2

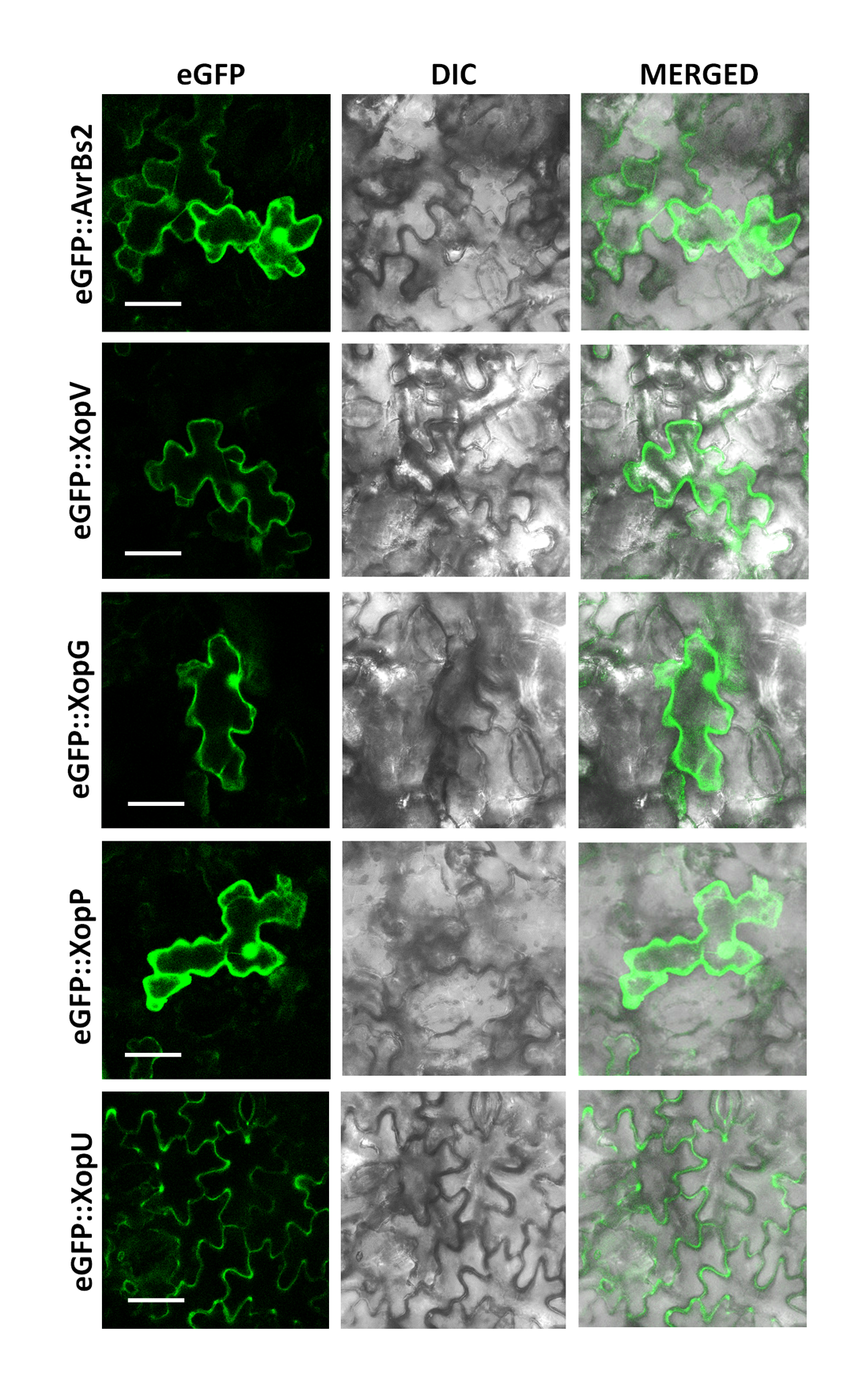

### Supplementary Fig S3

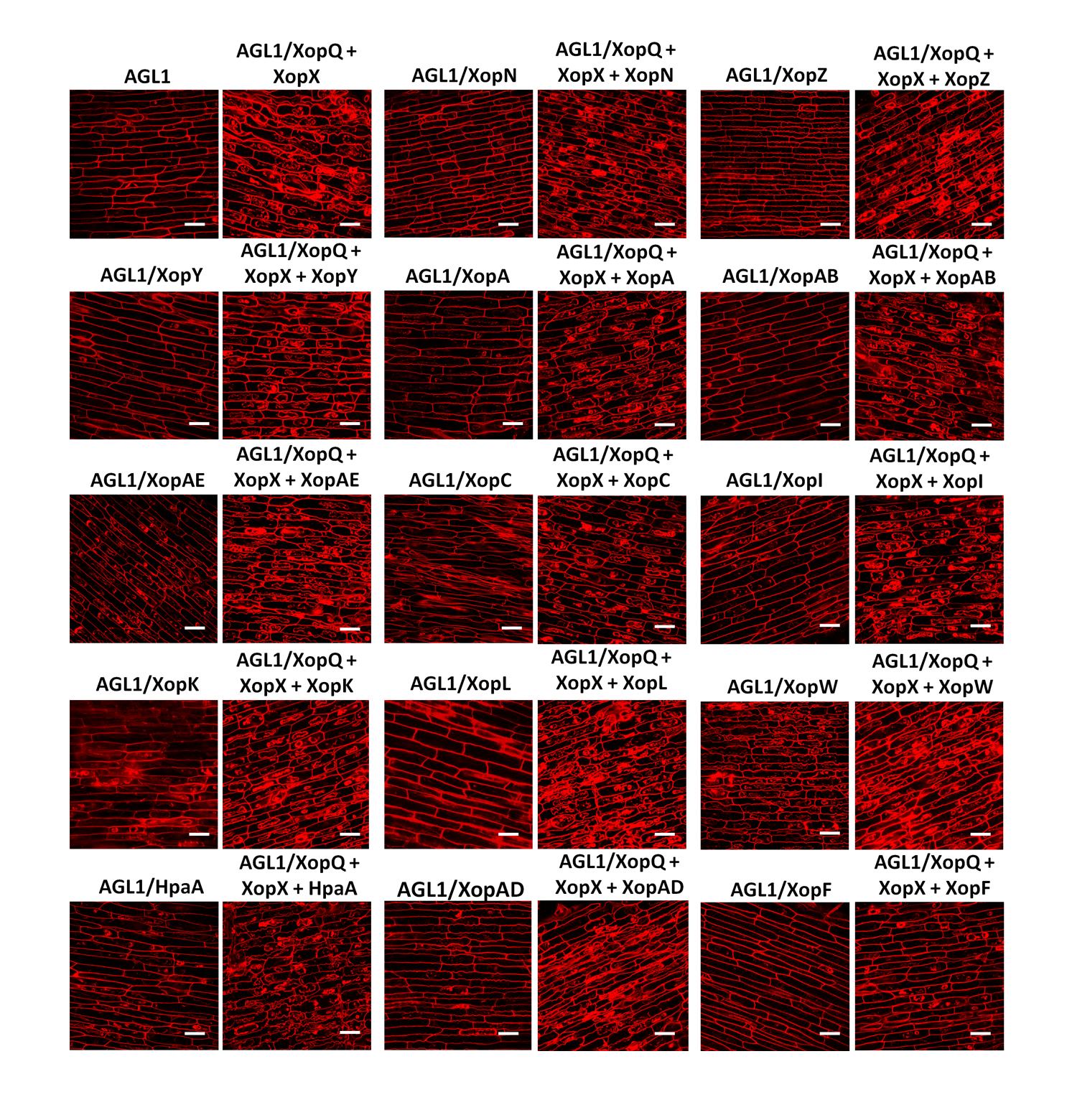

### Supplementary Fig S4

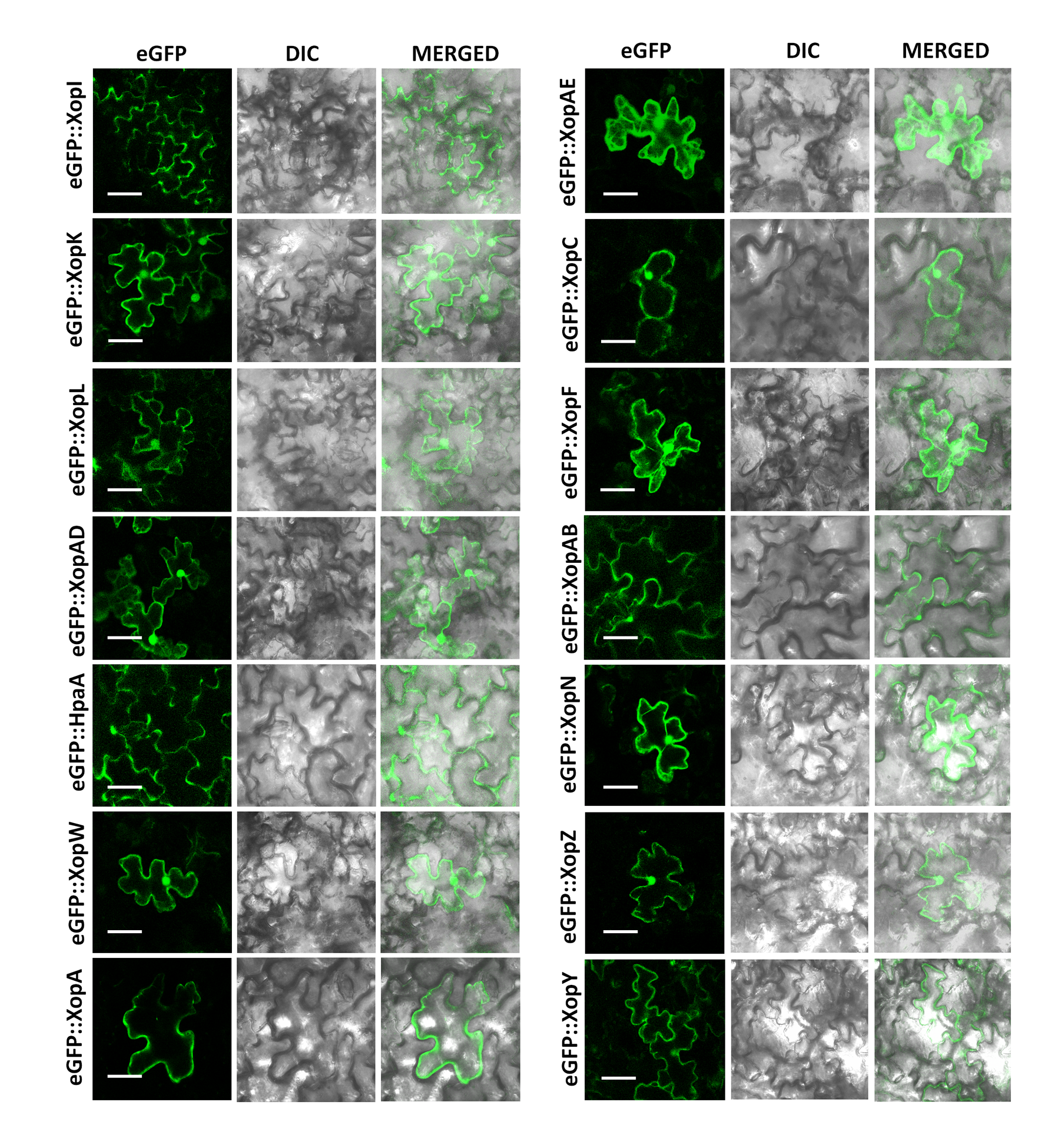

### Supplementary Fig S5

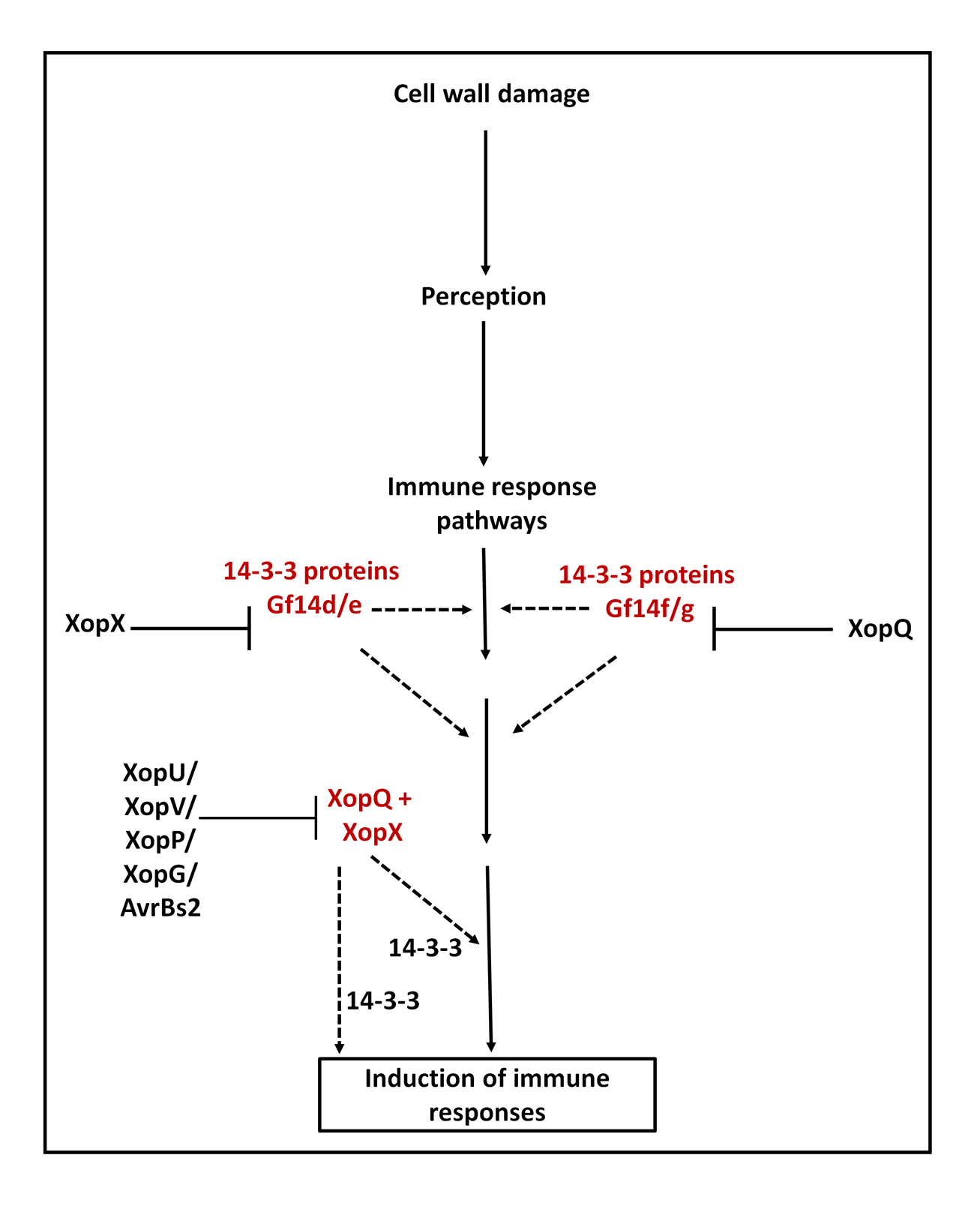
