## Supplementary table S1 for "Interaction of the *Xanthomonas* effectors XopQ and XopX results in induction of rice immune responses"

**Supplementary table 1. List of primers used in the study**

| **Primer name** | **Primer sequence** |
| --- | --- |
| M13F | GTAAAACGACGGCCAGT |
| M13R | GGAAACAGCTATGACCATG |
| *xopX* pENTR F | CACCATGCTGGAGTCCCAGGCGC |
| *xopX* pENTR R | TCAATGCAGCGTCGAAGGACGGTCGG |
| *xopX S84A* SDM F | ACCGCCGCTCCCGCCACTG |
| *xopX S84A* SDM R | AGCGGCGGTGCTTGCGCGTC |
| *xopX S193A* SDM F | ATTGCCAACCCTGAAAACGGACTGGA |
| *xopX S193A* SDM R | GTTGGCAATGGCCCCGCGGA |
| *xopX S193D* SDM F | ATTGACAACCCTGAAAACGGACTGGAAG |
| *xopX S193D* SDM R | GTTGTCAATGGCCCCGCGGAG |
| *xopX S430A* SDM F | CTTCGCCGGGCCGGGCCAG |
| *xopX S430A* SDM R | CCCGGCGAAGTCTCGGCGCAC |
| *xopX S477A* SDM F | GAGGCCATTCCACGGCAGCTATCTAACAAC |
| *xopX S477A* SDM R | GAATGGCCTCAGACCGGCCC |
| *xopX S477D* SDM F | GAGGACATTCCACGGCAGCTATCTAACAAC |
| *xopX S477D* SDM R | GAATGTCCTCAGACCGGCCCGG |
| *xopX T621A* SDM F | TTCGCCGGGCCCGACGGC |
| *xopX T621A* SDM R | CCCGGCGAACAGCCGGCT |
| *xopX* HindIII pHM1 F | TCCCAAGCTTATGCTGGAGTCCCAGGCGC |
| *xopX* EcoRI pHM1 R | TCCGGAATTCTCAATGATGATGATGATGATGATGCAGCGTCGAAGGACGGTCGGC |
