## Supplementary table S2 for "Interaction of the *Xanthomonas* effectors XopQ and XopX results in induction of rice immune responses"

**Supplementary table S2: List of plasmids used in the study**

| **Plasmids** | **Characteristics** | **References** |
| --- | --- | --- |
| **pENTR clones** | | |
| pENTR/D-TOPO | cloning vector; Km^r^ | Invitrogen |
| pENTR/D-TOPO::*xopX* | pENTR/D-TOPO with 2,189-bp fragment containing full length *xopX* gene from BXO43 genomic DNA; Km^r^ | This study |
| pENTR/D-TOPO::*xopX S84A* | pENTR/D-TOPO containing full length *xopX S84A* gene obtained by SDM from pENTR/D-TOPO::*xopX*; Km^r^ | This study |
| pENTR/D-TOPO::*xopX S193A* | pENTR/D-TOPO containing full length *xopX S193A* gene obtained by SDM from pENTR/D-TOPO::*xopX*; Km^r^ | This study |
| pENTR/D-TOPO::*xopX S193D* | pENTR/D-TOPO containing full length *xopX S193D* gene obtained by SDM from pENTR/D-TOPO::*xopX*; Km^r^ | This study |
| pENTR/D-TOPO::*xopX T430A* | pENTR/D-TOPO containing full length *xopX T430A* gene obtained by SDM from pENTR/D-TOPO::*xopX*; Km^r^ | This study |
| pENTR/D-TOPO::*xopX S477A* | pENTR/D-TOPO containing full length *xopX S477A* gene obtained by SDM from pENTR/D-TOPO::*xopX*; Km^r^ | This study |
| pENTR/D-TOPO::*xopX S477D* | pENTR/D-TOPO containing full length *xopX S477D* gene obtained by SDM from pENTR/D-TOPO::*xopX*; Km^r^ | This study |
| pENTR/D-TOPO::*xopX T621A* | pENTR/D-TOPO containing full length *xopX T621A* gene obtained by SDM from pENTR/D-TOPO::*xopX*; Km^r^ | This study |
| pENTR/D-TOPO::*avrBs2* | pENTR/D-TOPO with 2145-bp fragment containing full length *avrBs2* gene from BXO43; Km^r^ | This study |
| pENTR/D-TOPO::*xopC2* | pENTR/D-TOPO with 1791-bp fragment containing full length *xopC2* gene from BXO43; Km^r^ | This study |
| pENTR/D-TOPO::*xopF* | pENTR/D-TOPO with 1986-bp fragment containing full length *xopF* gene from BXO43; Km^r^ | This study |
| pENTR/D-TOPO::*xopG* | pENTR/D-TOPO with 468-bp fragment containing full length *xopG* gene from BXO43; Km^r^ | This study |
| pENTR/D-TOPO::*xopI* | pENTR/D-TOPO with 573-bp fragment containing full length *xopI* gene from BXO43; Km^r^ | This study |
| pENTR/D-TOPO::*xopK* | pENTR/D-TOPO with 2538-bp fragment containing full length *xopK* gene from BXO43; Km^r^ | This study |
| pENTR/D-TOPO::*xopL* | pENTR/D-TOPO with 1959-bp fragment containing full length *xopL* gene from BXO43; Km^r^ | This study |
| pENTR/D-TOPO::*xopP* | pENTR/D-TOPO with 2190-bp fragment containing full length *xopP* gene from BXO43; Km^r^ | This study |
| pENTR/D-TOPO::*xopU* | pENTR/D-TOPO with 2958-bp fragment containing full length *xopU* gene from BXO43; Km^r^ | This study |
| pENTR/D-TOPO::*xopV* | pENTR/D-TOPO with 996-bp fragment containing full length *xopV* gene from BXO43; Km^r^ | This study |
| pENTR/D-TOPO::*xopW* | pENTR/D-TOPO with 615-bp fragment containing full length *xopW* gene from BXO43; Km^r^ | This study |
| pENTR/D-TOPO::*xopY* | pENTR/D-TOPO with 846-bp fragment containing full length *xopY* gene from BXO43; Km^r^ | This study |
| pENTR/D-TOPO::*xopAB* | pENTR/D-TOPO with 582-bp fragment containing full length *xopAB* gene from BXO43; Km^r^ | This study |
| pENTR/D-TOPO::*xopAD* | pENTR/D-TOPO with 8784-bp fragment containing full length *xopAD* gene from BXO43; Km^r^ | This study |
| pENTR/D-TOPO::*xopAE* | pENTR/D-TOPO with 1941-bp fragment containing full length *xopAE* gene from BXO43; Km^r^ | This study |
| pENTR/D-TOPO::*xopA* | pENTR/D-TOPO with 420-bp fragment containing full length *xopA* gene from BXO43; Km^r^ | This study |
| pENTR/D-TOPO::*hpaA* | pENTR/D-TOPO with 828-bp fragment containing full length *hpaA* gene from BXO43; Km^r^ | This study |
| **pHM1 overexpression clones** | |  |
| pHM1 | Broad host range vector for overexpression of proteins in *X. oryzae* pv. *oryzae*; Sp^r^ | (Innes et al., 1988) |
| pHM1::*xopX* | pHM1 containing full length *xopX* gene from pENTR/D-TOPO::*xopX* cloned into HindIII and EcoRI sites of pHM1; Sp^r^ | This study |
| pHM1::*xopX S84A* | pHM1 containing full length *xopX S84A* gene from pENTR/D-TOPO:: *xopX S84A* cloned into HindIII and EcoRI sites of pHM1; Sp^r^ | This study |
| pHM1::*xopX S193A* | pHM1 containing full length *xopX S193A* gene from pENTR/D-TOPO:: *xopX S193A* cloned into HindIII and EcoRI sites of pHM1; Sp^r^ | This study |
| pHM1::*xopX S193D* | pHM1 containing full length *xopX S193D* gene from pENTR/D-TOPO:: *xopX S193D* cloned into HindIII and EcoRI sites of pHM1; Sp^r^ | This study |
| pHM1::*xopX T430A* | pHM1 containing full length *xopX T430A* gene from pENTR/D-TOPO::*xopX T430A* cloned into HindIII and EcoRI sites of pHM1; Sp^r^ | This study |
| pHM1::*xopX S477A* | pHM1 containing full length *xopX S477A* gene from pENTR/D-TOPO::*xopX S477A* cloned into HindIII and EcoRI sites of pHM1; Sp^r^ | This study |
| pHM1::*xopX S477D* | pHM1 containing full length *xopX S477D* gene from pENTR/D-TOPO::*xopX S477D* cloned into HindIII and EcoRI sites of pHM1; Sp^r^ | This study |
| pHM1::*xopX T621A* | pHM1 containing full length *xopX T621A* gene from pENTR/D-TOPO:: *xopX T621A* cloned into HindIII and EcoRI sites of pHM1; Sp^r^ | This study |
| **Plant overexpression clones** | |  |
| pH7WGF2 | Plant overexpression vector with constitutive 35S CaMV promoter and N-terminal eGFP fusion; Sp^r^ | (Karimi et al., 2002) |
| eGFP::*xopQ* | pH7WGF2 containing full length *xopQ* gene obtained from recombination of pENTR::*xopQ* with pH7WGF2; Sp^r^ | This study |
| eGFP::*xopX* | pH7WGF2 containing full length *xopX* gene obtained from recombination of pENTR::*xopX* with pH7WGF2; Sp^r^ | This study |
| eGFP::*xopX S193A* | pH7WGF2 containing full length *xopX S193A* gene obtained from recombination of pENTR::*xopX S193A* with pH7WGF2; Sp^r^ | This study |
| eGFP::*xopX S193D* | pH7WGF2 containing full length *xopX S193D* gene obtained from recombination of pENTR::*xopX S193D* with pH7WGF2; Sp^r^ | This study |
| eGFP::*xopX S477A* | pH7WGF2 containing full length *xopX S477A* gene obtained from recombination of pENTR::*xopX S477A* with pH7WGF2; Sp^r^ | This study |
| eGFP::*xopX S477D* | pH7WGF2 containing full length *xopX S477D* gene obtained from recombination of pENTR::*xopX S477D* with pH7WGF2; Sp^r^ | This study |
| eGFP::*avrBs2* | pH7WGF2 containing full length *avrBs2* gene obtained from recombination of pENTR:: *avrBs2* with pH7WGF2; Sp^r^ | This study |
| eGFP::*xopC2* | pH7WGF2 containing full length *xopC2* gene obtained from recombination of pENTR::*xopC2* with pH7WGF2; Sp^r^ | This study |
| eGFP::*xopF* | pH7WGF2 containing full length *xopF* gene obtained from recombination of pENTR::*xopF* with pH7WGF2; Sp^r^ | This study |
| eGFP::*xopG* | pH7WGF2 containing full length *xopG* gene obtained from recombination of pENTR::*xopG* with pH7WGF2; Sp^r^ | This study |
| eGFP::*xopI* | pH7WGF2 containing full length *xopI* gene obtained from recombination of pENTR::*xopI* with pH7WGF2; Sp^r^ | This study |
| eGFP::*xopK* | pH7WGF2 containing full length *xopK* gene obtained from recombination of pENTR::*xopK* with pH7WGF2; Sp^r^ | This study |
| eGFP::*xopL* | pH7WGF2 containing full length *xopL* gene obtained from recombination of pENTR::*xopL* with pH7WGF2; Sp^r^ | This study |
| eGFP:: *xopN* | pH7WGF2 containing full length *xopN* gene obtained from recombination of pENTR::*xopN* with pH7WGF2; Sp^r^ | This study |
| eGFP::*xopP* | pH7WGF2 containing full length *xopP* gene obtained from recombination of pENTR::*xopP* with pH7WGF2; Sp^r^ | This study |
| eGFP::*xopU* | pH7WGF2 containing full length *xopU* gene obtained from recombination of pENTR::*xopU* with pH7WGF2; Sp^r^ | This study |
| eGFP::*xopV* | pH7WGF2 containing full length *xopV* gene obtained from recombination of pENTR::*xopV* with pH7WGF2; Sp^r^ | This study |
| eGFP::*xopW* | pH7WGF2 containing full length *xopW* gene obtained from recombination of pENTR::*xopW* with pH7WGF2; Sp^r^ | This study |
| eGFP::*xopY* | pH7WGF2 containing full length *xopY* gene obtained from recombination of pENTR::*xopY* with pH7WGF2; Sp^r^ | This study |
| eGFP::*xopAB* | pH7WGF2 containing full length *xopAB* gene obtained from recombination of pENTR::*xopAB* with pH7WGF2; Sp^r^ | This study |
| eGFP::*xopAD* | pH7WGF2 containing full length *xopAD* gene obtained from recombination of pENTR::*xopAD* with pH7WGF2; Sp^r^ | This study |
| eGFP::*xopAE* | pH7WGF2 containing full length *xopAE* gene obtained from recombination of pENTR::*xopAE* with pH7WGF2; Sp^r^ | This study |
| eGFP::*xopA* | pH7WGF2 containing full length *xopA* gene obtained from recombination of pENTR::*xopA* with pH7WGF2; Sp^r^ | This study |
| eGFP::*hpaA* | pH7WGF2 containing full length *hpaA* gene obtained from recombination of pENTR::*hpaA* with pH7WGF2; Sp^r^ | This study |
| **BiFC clones** | |  |
| pDEST-VYCE(R)GW | Plant expression vector for BiFC having C-terminal portion of VFP as an N-terminal fusion to protein of interest; Km^r^ | (Gehl et al., 2009) |
| pDEST-VYNE(R)GW | Plant expression vector for BiFC having N-terminal portion of VFP as an N-terminal fusion to protein of interest; Km^r^ | (Gehl et al., 2009) |
| nVFP::*xopQ* | pDEST-VYNE(R)GW containing full length *xopQ* gene obtained from recombination of pENTR::*xopQ* with pDEST-VYCE(R)GW; Km^r^ | This study |
| cVFP::*xopX* | pDEST-VYCE(R)GW containing full length *xopX* gene obtained from recombination of pENTR::*xopX* with pDEST-VYCE(R)GW; Km^r^ | This study |
| cVFP::*xopX S193A* | pDEST-VYCE(R)GW containing full length *xopX S193A* gene obtained from recombination of pENTR::*xopX S193A* with pDEST-VYCE(R)GW; Km^r^ | This study |
| cVFP::*xopX S193D* | pDEST-VYCE(R)GW containing full length *xopX S193A* gene obtained from recombination of pENTR::*xopX S193D* with pDEST-VYCE(R)GW; Km^r^ | This study |
| cVFP::*xopX S477A* | pDEST-VYCE(R)GW containing full length *xopX S477A* gene obtained from recombination of pENTR::*xopX S477A* with pDEST-VYCE(R)GW; Km^r^ | This study |
| cVFP::*xopX S477D* | pDEST-VYCE(R)GW containing full length *xopX S477A* gene obtained from recombination of pENTR::*xopX S477D* with pDEST-VYCE(R)GW; Km^r^ | This study |
| nVFP::*gf14d* | pDEST-VYNE(R)GW containing full length *gf14d* gene obtained from recombination of pENTR::*gf14d* with pDEST-VYNE(R)GW; Km^r^ | (Deb et al., 2019) |
| nVFP::*gf14e* | pDEST-VYNE(R)GW with 789-bp fragment containing full length *gf14e* gene obtained from recombination of pENTR::*gf14e* with pDEST-VYNE(R)GW; Km^r^ | (Deb et al., 2019) |
| **Yeast expression clones** | | |
| pDEST32 | Expression vector for yeast 2-hybrid having the binding domain (BD) of a transcription factor as an N-terminal fusion to the gene of interest for activation of reporter genes; Gent^r^ | Invitrogen |
| pDEST22 | Expression vector for yeast 2-hybrid having the activation domain (AD) of a transcription factor as an N-terminal fusion to the gene of interest for activation of reporter genes; Amp^r^ | Invitrogen |
| AD::*xopQ* | pDEST32 containing full length *xopQ* gene obtained from recombination of pENTR::*xopQ* with pDEST22; Amp^r^ | This study |
| AD::*xopN* | pDEST32 containing full length *xopN* gene obtained from recombination of pENTR::*xopN* with pDEST22; Amp^r^ | This study |
| AD::*xopZ* | pDEST32 containing full length *xopZ* gene obtained from recombination of pENTR::*xopZ* with pDEST22; Amp^r^ | This study |
| BD::*xopX* | pDEST32 containing full length *xopX* gene obtained from recombination of pENTR::*xopX* with pDEST32; Gent^r^ | This study |
| BD::*xopX S84A* | pDEST32 containing full length *xopX S84A* gene obtained from recombination of pENTR::*xopX S84A* with pDEST32; Gent^r^ | This study |
| BD::*xopX S193A* | pDEST32 containing full length *xopX S193A* gene obtained from recombination of pENTR::*xopX S193A* with pDEST32; Gent^r^ | This study |
| BD::*xopX S193D* | pDEST32 containing full length *xopX S193D* gene obtained from recombination of pENTR::*xopX S193D* with pDEST32; Gent^r^ | This study |
| BD::*xopX T430A* | pDEST32 containing full length *xopX T430A* gene obtained from recombination of pENTR::*xopX T430A* with pDEST32; Gent^r^ | This study |
| BD::*xopX S477A* | pDEST32 containing full length *xopX S477A* gene obtained from recombination of pENTR::*xopX S477A* with pDEST32; Gent^r^ | This study |
| BD::*xopX S477D* | pDEST32 containing full length *xopX S477D* gene obtained from recombination of pENTR::*xopX S477D* with pDEST32; Gent^r^ | This study |
| BD::*xopX T621A* | pDEST32 containing full length *xopX T621A* gene obtained from recombination of pENTR::*xopX T621A* with pDEST32; Gent^r^ | This study |
| AD::*gf1a* | pDEST22 with 795-bp fragment containing full length *gf14a* gene obtained from recombination of pENTR::*gf14a* with pDEST22; Amp^r^ | (Deb et al., 2019) |
| AD::*gf14b* | pDEST22 with 789-bp fragment containing full length *gf14b* gene obtained from recombination of pENTR::*gf14b* with pDEST22; Amp^r^ | (Deb et al., 2019) |
| AD::*gf14c* | pDEST22 with 771-bp fragment containing full length *gf14c* gene obtained from recombination of pENTR::*gf14c* with pDEST22; Amp^r^ | (Deb et al., 2019) |
| AD::*gf14d* | pDEST22 with 798-bp fragment containing full length *gf14d* gene obtained from recombination of pENTR::*gf14d* with pDEST22; Amp^r^ | (Deb et al., 2019) |
| AD::*gf14e* | pDEST22 with 789-bp fragment containing full length *gf14e* gene obtained from recombination of pENTR::*gf14e* with pDEST22; Amp^r^ | (Deb et al., 2019) |
| AD::*gf14f* | pDEST22 with 783-bp fragment containing full length *gf14f* gene obtained from recombination of pENTR::*gf14f* with pDEST22; Amp^r^ | (Deb et al., 2019) |
| AD::*gf14g* | pDEST22 with 612-bp fragment containing full length *gf14g* gene obtained from recombination of pENTR::*gf14g* with pDEST22; Amp^r^ | (Deb et al., 2019) |
| AD::*gf14h* | pDEST22 with 693-bp fragment containing full length *gf14h* gene obtained from recombination of pENTR::*gf14h* with pDEST22; Amp^r^ | (Deb et al., 2019) |

DEB, S., GUPTA, M. K., PATEL, H. K. & SONTI, R. V. 2019. Xanthomonas oryzae pv. oryzae XopQ protein suppresses rice immune responses through interaction with two 14-3-3 proteins but its phospho-null mutant induces rice immune responses and interacts with another 14-3-3 protein. *Mol Plant Pathol*.

GEHL, C., WAADT, R., KUDLA, J., MENDEL, R. R. & HANSCH, R. 2009. New GATEWAY vectors for high throughput analyses of protein-protein interactions by bimolecular fluorescence complementation. *Mol Plant,* 2**,** 1051-8.

INNES, R. W., HIROSE, M. A. & KUEMPEL, P. L. 1988. Induction of nitrogen-fixing nodules on clover requires only 32 kilobase pairs of DNA from the Rhizobium trifolii symbiosis plasmid. *J Bacteriol,* 170**,** 3793-802.

KARIMI, M., INZE, D. & DEPICKER, A. 2002. GATEWAY vectors for Agrobacterium-mediated plant transformation. *Trends Plant Sci,* 7**,** 193-5.
