## Supplementary table S3 for "Interaction of the *Xanthomonas* effectors XopQ and XopX results in induction of rice immune responses"

**Supplementary table 3. List of strains used in the study**

| **Strains** | **Characteristics** | **References** |
| --- | --- | --- |
| ***Escherichia coli* strains** | |  |
| DH5α | λ^–^ f80dlacZDM15 D(lacZYA-argF) U169 recA1 endA hsdR17 (rK^–^ mK^–^) | Thermo Fischer Scientific |
| ***Xanthomonas oryzae* pv. *oryzae* strains** | | |
| BXO43 | *rif*-2; derivative of BXO1 (Wild- type; Indian isolate) | (Thieme et al., 2005) |
| QM | *xopN::*pK18mob *ΔxopQ ΔxopX ΔxopZ* quadruple mutant; *rif-2;* Km^r^ | (Sinha et al., 2013) |
| QM /pHM1 | *xopN::*pK18mob *ΔxopQ ΔxopX ΔxopZ; rif-2;* Km^r^ ; Sp^r^ ; derivative of QM | This study |
| QM/pHM1::*xopX* | *xopN::* pK18mob *ΔxopQ ΔxopX ΔxopZ; rif-2;* Km^r^ ; Sp^r^ ; XopX+, derivative of QM | This study |
| QM/pHM1::*xopX S84A* | *xopN::*pK18mob *ΔxopQ ΔxopX ΔxopZ; rif-2;* Km^r^ ; Sp^r^ ; XopX S84A, derivative of QM | This study |
| QM/pHM1::*xopX S193A* | *xopN::*pK18mob *ΔxopQ ΔxopX ΔxopZ; rif-2;* Km^r^ ; Sp^r^ ; XopX S193A, derivative of QM | This study |
| QM/pHM1::*xopX S193D* | *xopN::*pK18mob *ΔxopQ ΔxopX ΔxopZ; rif-2;* Km^r^ ; Sp^r^ ; XopX S193D, derivative of QM | This study |
| QM/pHM1::*xopX T430A* | *xopN::*pK18mob *ΔxopQ ΔxopX ΔxopZ; rif-2;* Km^r^ ; Sp^r^ ; XopX T430A, derivative of QM | This study |
| QM/pHM1::*xopX S477A* | *xopN::*pK18mob *ΔxopQ ΔxopX ΔxopZ; rif-2;* Km^r^ ; Sp^r^ ; XopX S477A, derivative of QM | This study |
| QM/pHM1::*xopX S477D* | *xopN::*pK18mob *ΔxopQ ΔxopX ΔxopZ; rif-2;* Km^r^ ; Sp^r^ ; XopX S477D, derivative of QM | This study |
| QM/pHM1::*xopX T621A* | *xopN::*pK18mob *ΔxopQ ΔxopX ΔxopZ; rif-2;* Km^r^ ; Sp^r^ ; XopX T621A, derivative of QM | This study |
| ***Agrobacterium tumefaciens strains*** | | |
| AGL1 | AGL0 (*C58 pTiBo542*) *recA*::*bla*, T-region deleted Mop(+) Cb(R) (AGL0 is an EHA101 with the T-region deleted) | (Lazo et al., 1991) |
| AGL1/eGFP::*xopQ* | AGL1 harboring plasmid pH7WGF2::*xopQ*; Sp^r^ | This study |
| AGL1/eGFP::*xopX* | AGL1 harboring plasmid pH7WGF2::*xopX*; Sp^r^ | This study |
| AGL1/eGFP::*xopX S193A* | AGL1 harboring plasmid  pH7WGF2::*xopX S193A*; Sp^r^ | This study |
| AGL1/eGFP::*xopX S193D* | AGL1 harboring plasmid  pH7WGF2::*xopX S193D*; Sp^r^ | This study |
| AGL1/eGFP::*xopX*  *S477A* | AGL1 harboring plasmid  pH7WGF2::*xopX S477A*; Sp^r^ | This study |
| AGL1/eGFP::*xopX*  *S477D* | AGL1 harboring plasmid  pH7WGF2::*xopX S477D*; Sp^r^ | This study |
| AGL1/ eGFP::*avrBs2* | AGL1 harboring plasmid pH7WGF2::*avrBs2*; Sp^r^ | This study |
| AGL1/ eGFP::*xopC* | AGL1 harboring plasmid pH7WGF2::*xopC*; Sp^r^ | This study |
| AGL1/ eGFP::*xopF* | AGL1 harboring plasmid pH7WGF2::*xopF*; Sp^r^ | This study |
| AGL1/ eGFP::*xopG* | AGL1 harboring plasmid pH7WGF2::*xopG*; Sp^r^ | This study |
| AGL1/ eGFP::*xopI* | AGL1 harboring plasmid pH7WGF2::*xopI*; Sp^r^ | This study |
| AGL1/ eGFP::*xopK* | AGL1 harboring plasmid pH7WGF2::*xopK*; Sp^r^ | This study |
| AGL1/ eGFP::*xopL* | AGL1 harboring plasmid pH7WGF2::*xopL*; Sp^r^ | This study |
| AGL1/ eGFP::*xopN* | AGL1 harboring plasmid pH7WGF2::*xopN*; Sp^r^ | This study |
| AGL1/ eGFP::*xopP* | AGL1 harboring plasmid pH7WGF2::*xopP*; Sp^r^ | This study |
| AGL1/ eGFP::*xopU* | AGL1 harboring plasmid pH7WGF2::*xopU*; Sp^r^ | This study |
| AGL1/ eGFP::*xopV* | AGL1 harboring plasmid pH7WGF2::*xopV*; Sp^r^ | This study |
| AGL1/eGFP::*xopW* | AGL1 harboring plasmid pH7WGF2::*xopW*; Sp^r^ | This study |
| AGL1/ eGFP::*xopY* | AGL1 harboring plasmid pH7WGF2::*xopY*; Sp^r^ | This study |
| AGL1/eGFP::*xopAB* | AGL1 harboring plasmid pH7WGF2::*xopAB*; Sp^r^ | This study |
| AGL1/ eGFP::*xopAD* | AGL1 harboring plasmid pH7WGF2::*xopAD*; Sp^r^ | This study |
| AGL1/ eGFP:: *xopAE* | AGL1 harboring plasmid pH7WGF2::*xopAE*; Sp^r^ | This study |
| AGL1/ eGFP::*xopA* | AGL1 harboring plasmid pH7WGF2::*xopA*; Sp^r^ | This study |
| AGL1/ eGFP::*hpaA* | AGL1 harboring plasmid pH7WGF2::*hpaA*; Sp^r^ | This study |
| AGL1/pDEST-VYCE(R)GW | AGL1 harboring plasmid pDEST-VYCE(R)GW; Km^r^ | This study |
| AGL1/pDEST-VYNE(R)GW | AGL1 harboring plasmid pDEST-VYNE(R)GW; Km^r^ | This study |
| AGL1/nVFP::*xopQ* | AGL1 harboring plasmid pDEST-VYNE(R)GW::*xopQ*; Km^r^ | (Deb et al., 2019) |
| AGL1/cVFP::*xopX* | AGL1 harboring plasmid pDEST-VYCE(R)GW::*xopX*; Km^r^ | This study |
| AGL1/cVFP::*xopX S193A* | AGL1 harboring plasmid pDEST-VYCE(R)GW::*xopX S193A*; Km^r^ | This study |
| AGL1/cVFP::*xopX S193D* | AGL1 harboring plasmid pDEST-VYCE(R)GW::*xopX S193D*; Km^r^ | This study |
| AGL1/cVFP::*xopX S477A* | AGL1 harboring plasmid pDEST-VYCE(R)GW::*xopX S477A*; Km^r^ | This study |
| AGL1/cVFP::*xopX S477D* | AGL1 harboring plasmid pDEST-VYCE(R)GW::*xopX S477D*; Km^r^ | This study |
| AGL1/nVFP::*gf14d* | AGL1 harboring plasmid pDEST-VYNE(R)GW::*gf14d*; Km^r^ | This study |
| AGL1/nVFP::*gf14e* | AGL1 harboring plasmid pDEST-VYNE(R)GW::*gf14e*; Km^r^ | (Deb et al., 2019) |
| ***Saccharomyces cerevisiae strains*** | |  |
| pJ694a | MATa trp1-901 leu2-3,112 ura3-52 his3-200 gal4(deleted) gal80(deleted) LYS2::GAL1-HIS3 GAL2-ADE2 met2::GAL7-lacZ | (James et al., 1996) |
| pJ694a/pDEST32 | pJ694a harboring plasmid pDEST32;  -LEU | This study |
| pJ694a/pDEST22 | pJ694a harboring plasmid pDEST22;  -TRP | This study |
| pJ694a/AD::*xopQ* | pJ694a harboring plasmid pDEST22::*xopQ;* -TRP | (Deb et al., 2019) |
| pJ694a/AD::*xopN* | pJ694a harboring plasmid pDEST22::*xopN;* -TRP | This study |
| pJ694a/AD::*xopZ* | pJ694a harboring plasmid pDEST22::*xopZ;* -TRP | This study |
| pJ694a/BD::*xopX* | pJ694a harboring plasmid pDEST32::*xopX;* -LEU | This study |
| pJ694a/BD::*xopX S84A* | pJ694a harboring plasmid pDEST32::*xopX S84A;* -LEU | This study |
| pJ694a/BD::*xopX S193A* | pJ694a harboring plasmid pDEST32::*xopX S193A;* -LEU | This study |
| pJ694a/BD::*xopX S193D* | pJ694a harboring plasmid pDEST32::*xopX S193D;* -LEU | This study |
| pJ694a/BD::*xopX T430A* | pJ694a harboring plasmid pDEST32::*xopX T430A;* -LEU | This study |
| pJ694a/BD::*xopX S477A* | pJ694a harboring plasmid pDEST32::*xopX S477A;* -LEU | This study |
| pJ694a/BD::*xopX S477D* | pJ694a harboring plasmid pDEST32::*xopX S477D;* -LEU | This study |
| pJ694a/BD::*xopX T621A* | pJ694a harboring plasmid pDEST32::*xopX T621A;* -LEU | This study |
| pJ694a/AD::*gf14a* | pJ694a harboring plasmid pDEST22::*gf14a; -*TRP | (Deb et al., 2019) |
| pJ694a/AD::*gf14b* | pJ694a harboring plasmid pDEST22::*gf14b;* -TRP | (Deb et al., 2019) |
| pJ694a/AD::*gf14c* | pJ694a harboring plasmid pDEST22::*gf14c; -*TRP | (Deb et al., 2019) |
| pJ694a/AD::*gf14d* | pJ694a harboring plasmid pDEST22::*gf14d; -*TRP | (Deb et al., 2019) |
| pJ694a/AD::*gf14e* | pJ694a harboring plasmid pDEST22::*gf14e; -*TRP | (Deb et al., 2019) |
| pJ694a/AD::*gf14f* | pJ694a harboring plasmid pDEST22::*gf14f; -*TRP | (Deb et al., 2019) |
| pJ694a/AD::*gf14g* | pJ694a harboring plasmid pDEST22::*gf14g; -*TRP | (Deb et al., 2019) |
| pJ694a/AD::*gf14h* | pJ694a harboring plasmid pDEST22::*gf14h; -*TRP | (Deb et al., 2019) |

DEB, S., GUPTA, M. K., PATEL, H. K. & SONTI, R. V. 2019. Xanthomonas oryzae pv. oryzae XopQ protein suppresses rice immune responses through interaction with two 14-3-3 proteins but its phospho-null mutant induces rice immune responses and interacts with another 14-3-3 protein. *Mol Plant Pathol*.

JAMES, P., HALLADAY, J. & CRAIG, E. A. 1996. Genomic libraries and a host strain designed for highly efficient two-hybrid selection in yeast. *Genetics,* 144**,** 1425-36.

LAZO, G. R., STEIN, P. A. & LUDWIG, R. A. 1991. A DNA transformation-competent Arabidopsis genomic library in Agrobacterium. *Biotechnology (N Y),* 9**,** 963-7.

SINHA, D., GUPTA, M. K., PATEL, H. K., RANJAN, A. & SONTI, R. V. 2013. Cell wall degrading enzyme induced rice innate immune responses are suppressed by the type 3 secretion system effectors XopN, XopQ, XopX and XopZ of Xanthomonas oryzae pv. oryzae. *PLoS One,* 8**,** e75867.

THIEME, F., KOEBNIK, R., BEKEL, T., BERGER, C., BOCH, J., BUTTNER, D., CALDANA, C., GAIGALAT, L., GOESMANN, A., KAY, S., KIRCHNER, O., LANZ, C., LINKE, B., MCHARDY, A. C., MEYER, F., MITTENHUBER, G., NIES, D. H., NIESBACH-KLOSGEN, U., PATSCHKOWSKI, T., RUCKERT, C., RUPP, O., SCHNEIKER, S., SCHUSTER, S. C., VORHOLTER, F. J., WEBER, E., PUHLER, A., BONAS, U., BARTELS, D. & KAISER, O. 2005. Insights into genome plasticity and pathogenicity of the plant pathogenic bacterium Xanthomonas campestris pv. vesicatoria revealed by the complete genome sequence. *J Bacteriol,* 187**,** 7254-66.
