## Supplementary table S4 for "Interaction of the *Xanthomonas* effectors XopQ and XopX results in induction of rice immune responses"

**Supplementary table 4. Details of annotated *X. oryzae pv. oryzae* effectors screened for suppression of XopQ- XopX mediated immune responses.**

| **S. No.** | **Genes encoding effectors** | **Gene ID** | **CDS Length** |
| --- | --- | --- | --- |
| 1 | *avrBs2* | XOO 0148 | 2145 bp |
| 2 | *xopC2* | XOO 3221 | 1791 bp |
| 3 | *xopF* | XOO 0103 | 1986 bp |
| 4 | *xopG* | XOO 4258 | 468 bp |
| 5 | *xopI* | XOO 3626 | 573 bp |
| 6 | *xopK* | XOO 1669 | 2538 bp |
| 7 | *xopL* | XOO 1662 | 1959 bp |
| 8 | *xopN* | XOO 0315 | 2162 bp |
| 9 | *xopP* | XOO 3222 | 2190 bp |
| 10 | *xopU* | XOO 2877 | 2958 bp |
| 11 | *xopV* | XOO 3803 | 996 bp |
| 12 | *xopW* | XOO 0037 | 615 bp |
| 13 | *xopY* | XOO 1488 | 846 bp |
| 14 | *xopZ* | XOO 2402 | 3866 bp |
| 15 | *xopAB* | XOO 3150 | 582 bp |
| 16 | *xopAD* | XOO 4145 | 8784 bp |
| 17 | *xopAE* | XOO 0110 | 1941 bp |
| 18 | *xopA* | XOO 0081 | 420 bp |
| 19 | *hpaA** | XOO 0097 | 828 bp |

* Type III secretion control protein, maybe not a type III effector (xanthomonas.org)
